# Structural mechanism of synergistic targeting of the *CX3CR1* nucleosome by PU.1 and C/EBP⍺

**DOI:** 10.1101/2023.08.25.554718

**Authors:** Tengfei Lian, Ruifang Guan, Bing-Rui Zhou, Yawen Bai

**Author notes:** These authors equally contributed to this work (Tengfei Lian, Ruifang Guan). Correspondence to: Yawen Bai; Tengfei Lian.

## Abstract

Pioneer transcription factors are vital for cell fate changes. PU.1 and C/EBP⍺ work together to regulate hematopoietic stem cell differentiation. However, how they recognize *in vivo* nucleosomal DNA targets remain elusive. Here we report the structures of the nucleosome containing the mouse genomic *CX3CR1* enhancer DNA and its complexes with PU.1 alone and with both PU.1 and the C/EBP⍺ DNA binding domain. Our structures reveal that PU.1 binds the DNA motif at the exit linker, shifting 17 bp of DNA into the core region through interactions with H2A, unwrapping ∼20 bp of nucleosomal DNA. C/EBP⍺ binding, aided by PU.1’s repositioning, unwraps ∼25 bp entry DNA. The PU.1 Q218H mutation, linked to acute myeloid leukemia, disrupts PU.1-H2A interactions. PU.1 and C/EBP⍺ jointly displace linker histone H1 and open the H1-condensed nucleosome array. Our study unveils how two pioneer factors can work cooperatively to open closed chromatin by altering DNA positioning in the nucleosome.

## Main

Gene expression and its regulation hold pivotal significance in governing cell fate and reprogramming processes.^1,2^ Nonetheless, within eukaryotic cells, the genomic DNA organizes itself into chromatin via the association of core and linker histones, forming nucleosomes and chromatosomes as primary structural units.^3,4^ In conjunction with the assembly of chromatin into more compact structures, a substantial portion of the DNA surface remains inaccessible to transcription factors. Pioneering transcription factors constitute a distinctive category of proteins capable of identifying closed chromatin regions, which are resistant to DNase I.^5,6^ These factors exhibit the ability to locally open chromatin, facilitating subsequent recruitment of other transcription factors. This coordination contributes to the regulation of gene expression, often assuming indispensable roles in processes such as cellular differentiation and development.

Although the structural basis of a transcription factor recognition of its cognitive motif in free DNA is well understood, how it targets the nucleosomal DNA in chromatin remains elusive. Current structural studies primarily use nucleosomes containing non-genomic DNA to form stable structures.^7,8,9,10,11,12^ The DNA motifs for binding the transcription factors are incorporated into them by either design or *in vitro* selection from a DNA library, leading to various transcription factor binding modes. The structural basis of how transcription factors bind to the nucleosomes containing the *in vivo* DNA targets has only been investigated recently, but no structures of nucleosome-pioneer factor complexes have been determined.^13–15^ This is because the nucleosomes containing genomic DNA are often more fragile and tend to dissociate during sample preparation for structural studies using the single-particle cryo-EM method.

PU.1 (encoded by the *spi1* gene) is a master regulator for hematopoietic stem cell differentiation and is essential for the terminal differentiation of macrophages.^16^ PU.1 mutation and reduced expression have been associated with acute myeloid leukemia (AML) and agammaglobulinemia.^17,18,19^ PU.1 binds to its DNA motif alone or co-binds with other factors, such as the C/EBPα family nearby at the enhancers of cell type-specific genes, and regulates their expression.^20,21^ It can target compact chromatin and enhance chromatin accessibility.^22^ In contrast, C/EBP⍺ binding to nucleosome-enriched regions in chromatin is more dependent on PU.1, and knockdown of PU.1 results in a decreased C/EBP⍺ binding to de novo enhancers during B cell transdifferentiation.^23^ Genomic and chromatin state analyses of PU.1 and C/EBP⍺ targeting in B cells and macrophages have revealed that they bind nucleosome-enriched chromatin.^24^ Comparison of fibroblast nucleosome occupancy by MNase-seq signals and transcription factor binding by ChIP-seq in macrophages shows that the CX3 chemokine receptor 1 (*CX3CR1*) locus consists of a nucleosome targeted by both PU.1 and C/EBP⍺.^25^ In addition, previous *in vitro* biochemical studies have confirmed the binding of the nucleosome consisting of the 162 bp *CX3CR1* DNA by the pioneer factors.^25^ Fig. 1**a** illustrates the DNA sequence.

**Fig 1.**
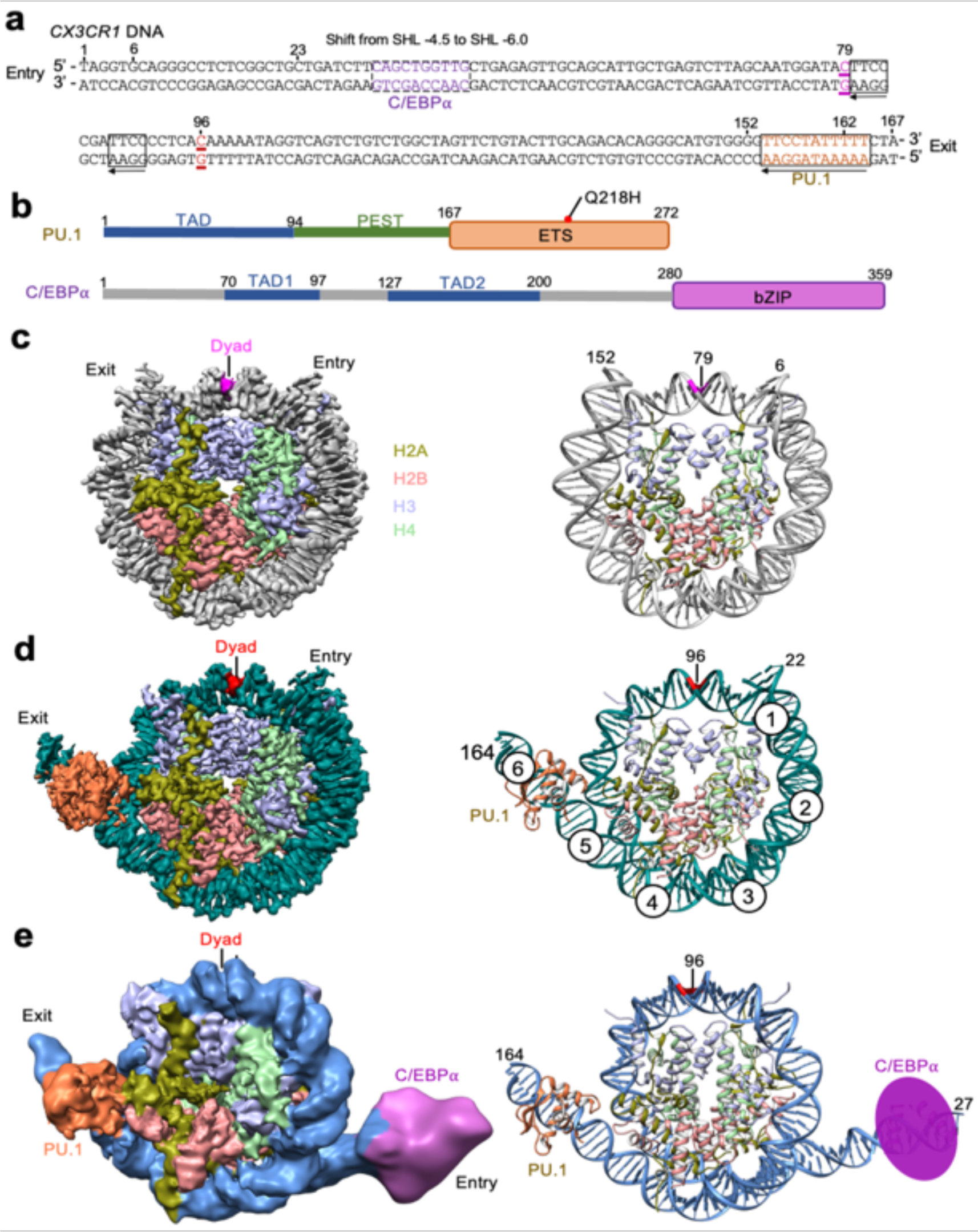
Structures of the *CX3CR1* nucleosome and its complexes with PU.1 and C/EBP⍺. **a.** Sequence of the *CX3CR1* DNA. PU.1 and C/EBP**⍺** DNA motifs are shown in the boxes. The arrows indicate the direction of the DNA motifs. Dyad positions in the free nucleosome and those bound to the transcription factors are underlined and in purple and red colors, respectively. **b.** Schematics of the PU.1 and C/EBP⍺ domain organization. Red spots indicate the mutations found in acute myeloid leukemia patients. **c-e.** Cryo-EM density maps (left) and the corresponding structural models (right) in ribbon for the free *CX3CR1* nucleosome, the nucleosome-PU.1 complex, and the nucleosome-PU.1-^DBD^C/EBP⍺ complex, respectively. For the structures of the nucleosome bound to PU.1 (**d**) and PU.1-^DBD^C/EBP⍺ (**e**), we used the disulfide mutant and 167 bp DNA. The numbers indicate the nucleotide number in the *CX3CR1* DNA.

PU.1 is a member of the erythroblast transformation-specific transcription factor family with the DNA binding domain (DBD or ETS) at its C-terminal region (Fig. 1**b**). In comparison, C/EBP⍺ is a transcription factor with a bZIP DNA binding domain at the C-terminal region and binds the DNA motif as a dimer (Fig. 1**b**). The N-terminal regions of both proteins are intrinsically disordered that are involved in interactions with other factors.^26,27^ The N-terminal region of PU.1 harbors a transactivation domain (TAD) that contains three acidic regions and a glutamate-rich region, and the N-terminal region of C/EBP⍺ contains two TAD domains. Both the N-terminal regions of PU.1 and C/EBP⍺ interact with basal transcription factors such as TFIID, TFIIB, and TBP,^28, 29^ and histone acetyltransferases CREB-binding protein (CBP)^30,31^ as well as SWI/SNF chromatin remodelers.^29,32^

In this study, we used cryo-EM and biochemical methods to investigate how PU.1 and C/EBP⍺ recognize the nucleosome containing the genomic *CX3CR1* enhancer DNA,^25^ the *in vivo* target of the transcription factors. Our study shows that PU.1 and C/EBP⍺ each can unwrap the nucleosomal DNA at the exit and entry sides, respectively. However, PU.1 binds the nucleosome with higher affinity by interacting with the core histone, leading to nucleosome repositioning, which shifts the binding site of C/EBP⍺ from the inner region of the nucleosome to the end region, facilitating the binding of C/EBP⍺. Our results reveal how two pioneer factors can synergistically bind to the nucleosome, unwrap nucleosomal DNA, evict linker histone H1, and open the H1-condensed nucleosome array.

## Results

### Structures of the free and pioneer factor-bound nucleosomes

To elucidate how PU.1 and C/EBP⍺ target the *CX3CR1* nucleosome, we reconstituted the nucleosome consisting of human core histones and the 162 bp DNA from the *CX3CR1* enhancer loci.^25^ We expressed and purified the recombinant full-length PU.1 and the DNA binding domain of C/EBP⍺ fused to maltose binding protein at the N-terminus to increase its solubility (Extended Data Fig. 1**a**). EMSA experiments showed that PU.1 binds the nucleosome with an apparent K_d_^app^ of ∼0.65 μM obtained by fitting the data to the Hill equation (Extended Data Fig. 1). In contrast, ^MBP-DBD^C/EBP⍺ binds the nucleosome with a lower affinity (K_d_^app^ of ∼1.7μM). Using the single-chain antibody (scFv)-assisted cryo-EM method,^33^ we obtained the density map of the nucleosome-scFv_2_ complex at an overall resolution of 2.6 Å (Fig. 1**c**, Extended Data Figs. 2, and Table 1). We also solved the structure of the nucleosome-scFv_2_-PU.1 complex at an overall resolution of 2.9 Å (Extended Data Figs. 3 and Table 1). However, we could not obtain a density map for the nucleosome-scFv_2_-^MBP-DBD^C/EBP⍺ complex, possibly due to its lower affinity. To investigate whether scFv affects the nucleosome binding by PU.1 and ^MBP-DBD^C/EBP⍺, we conducted EMSA experiments, showing that scFv had little effect on the apparent binding affinity of PU.1 and ^MBP-DBD^C/EBP⍺ to the nucleosome (Extended Data Fig. 1**g, h**).

**Table 1.**
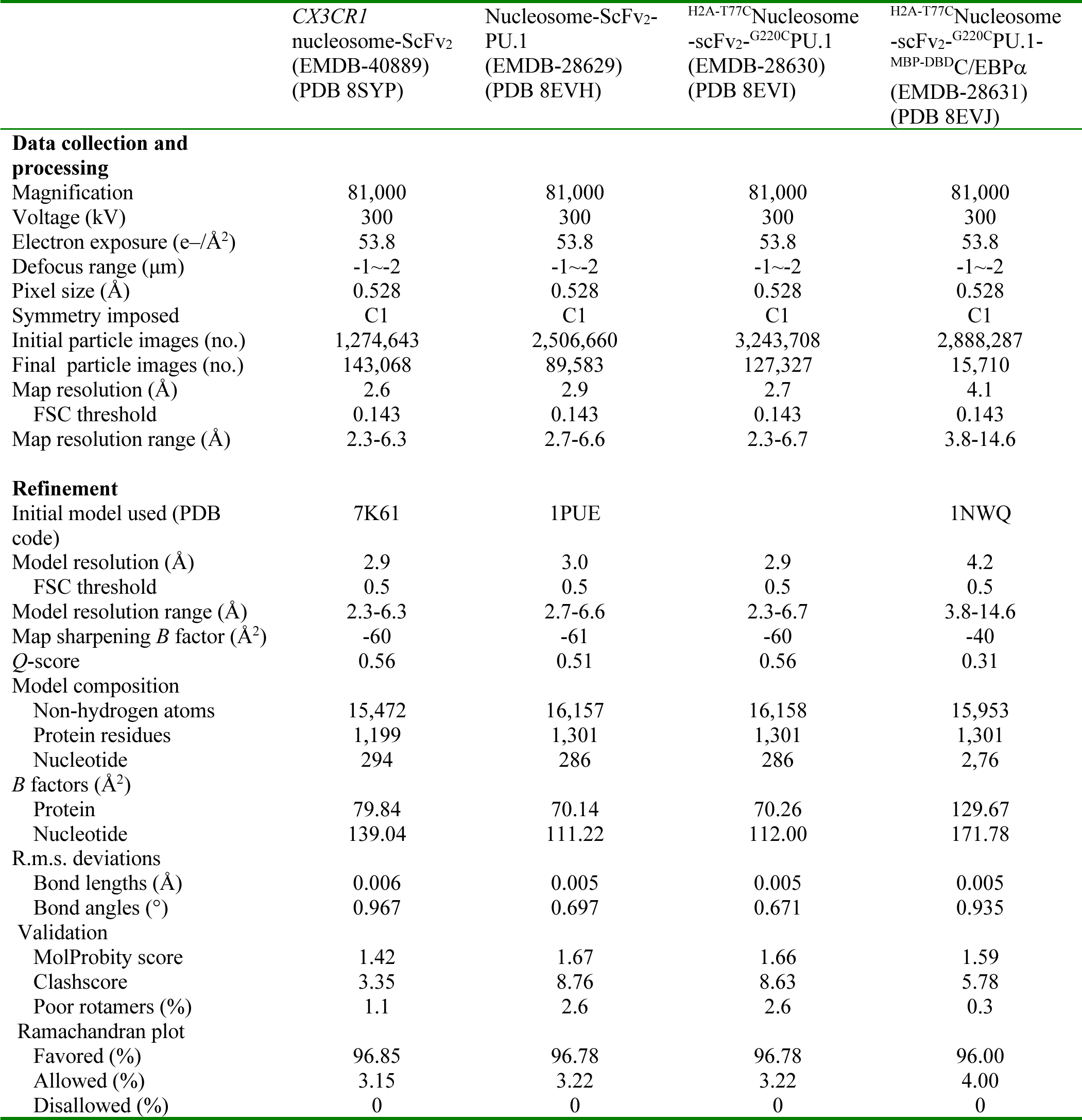
Cryo-EM data collection, refinement, and validation statistics.

The high-resolution density maps for the free *CX3CR1* nucleosome and its complex with PU.1 allow us to define the DNA base pairs unanimously (Fig. 2**a, b**), showing that the nucleosomes are uniquely positioned with nucleotides 79 and 96 at the dyad (Fig. 1**c, d**), respectively. Thus, PU.1 binding repositions the nucleosome by 17 bp. However, the density map for the PU.1 region has a local resolution of ∼6 Å (Extended Data Fig. 3), likely due to its dynamic motion. Nevertheless, the crystal structure of the DNA binding domain (DBD or ETS) of PU.1 bound to the DNA fragment (PDB ID: 1PUE) can fit the density map well, leading to a structure model showing that PU.1 G220 is close to H2A T77. To better define the structure, we engineered a disulfide bond by mutating these two residues to Cys to restrict the dynamic motion and used a longer DNA (167 bp, Fig. 1**a**) for PU.1 binding, which improved density map to an overall resolution of 2.7 Å and ∼4 Å for the PU.1 ETS binding region, respectively (Figure 1**d** and Extended Data Fig. 4). The new map shows densities for all core histones and backbone structures of the PU.1 ETS and residues interacting with DNA motif (Figure 1**d**, Extended Data Fig. 5**a-c**). The structural model derived from the density map of the mutated proteins fits the density of the wild-type (WT) complex very well (Extended Data Fig. 5**d**), indicating that the disulfide bond only restricts its dynamics without altering the structure of the complex.

**Fig. 2.**
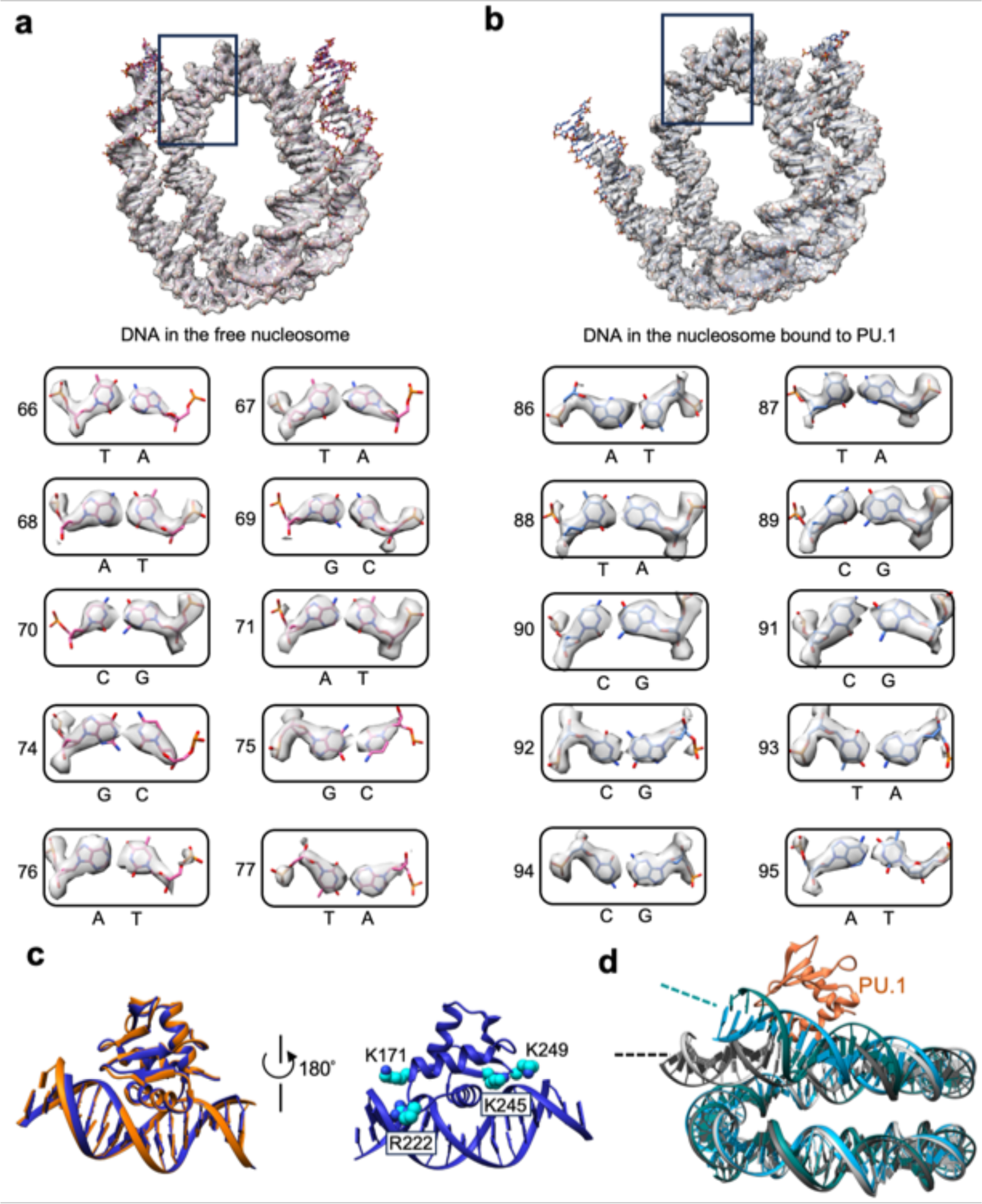
DNA registration assignment and modeling of PU.1 ETS binding to the nucleosome. **a.** Illustration of the assignment of representative DNA bases in the 162 bp free nucleosome. **b.** Illustration of the assignment of representative DNA bases in the 162 bp nucleosome-PU.1 (wild type) complex. **c.** Overlay of PU.1 ETS bound to the free DNA fragment (blue) and the unwrapped nucleosomal DNA (orange). Illustration of the interactions between positively charged residues (cyan) in the loops of PU.1 ETS and the phosphate backbone of DNA. **d**. Overlay of the free nucleosome (gray) and the nucleosome bound to PU.1 ETS (dark cyan), showing that PU.1 ETS bends the DNA towards it.

Furthermore, we obtain the density map for the nucleosome-scFv_2_-PU.1-^MBP-DBD^C/EBP⍺ complex with the disulfide bond and 167 bp DNA at 4.1 Å resolution (Figure 1**e**, Extended Data Fig. 6 and Table 1). The density map can be well fitted with the structural model of the nucleosome bound to the PU.1 ETS, except that the unwrapped ∼25 bp DNA on the entry side of the nucleosome. The unwrapped DNA is flexible and classified into three distinct conformations anchored around super-helical location (SHL) -4 (Extended Data Fig. 6). There are additional densities beyond the DNA at similar positions in all three conformations that can be fitted with the DNA binding region in the crystal structure of C/EBP⍺ bound to the DNA fragment (PDB: 1NWQ) (Extended Data Fig. 6**c** and h). This result suggests that the C/EBP⍺ dimer binds a region including a non-canonical motif in the unwrapped DNA at the entry side (corresponding to SHL -6.0) (Fig. 1**a**).

To confirm the binding of C/EBP⍺ at this location, we introduced a mutation in the associated DNA region, changing it from CAGC<u>TGG</u>TTG to CAGC<u>AAC</u>TTG (highlighted in the dashed box in Fig. 1**a**). The mutation resulted in a decrease in the apparent binding affinity between C/EBP⍺ and the nucleosome, with the K_d_^app^ value changing from 1.7 to 2.4 μM (Extended Data Fig. 7**a, b**). To further validate that the observed density on the DNA originated from C/EBP⍺, we incorporated the CAGC<u>TGG</u>TTG sequence to the ‘601’ nucleosome at the location corresponding to the one found in the *CX3CR1* nucleosome bound to PU.1 and C/EBP⍺ (Extended Data Fig. 7**d**). EMSA experiments demonstrated that C/EBP⍺ bound the chimeric nucleosome better than to the ‘601’ nucleosome (Extended Data Fig. 7**c, d**). Furthermore, we discovered that C/EBP⍺ also exhibited binding to a 20 bp double-stranded DNA fragment containing the CAGC<u>TGG</u>TTG sequence in the middle, while showing little binding to the fragment with the CAGC<u>AAC</u>TTG mutation (Extended Data Fig. 7**e**). Importantly, we established that MBP did not bind the free *CX3CR1* DNA and the *CX3CR1* nucleosome (Extended Data Fig. 7**f, g**), ruling out the possibility that the observed extra density originated from MBP. Based on our structural findings, we propose that PU.1 binds at the exit site and C/EBP⍺ binds at the entry site of the CX3CR1 nucleosome, which agrees with the ChIP-seq and MNase-seq results.^25^

### Interactions between PU.1 and nucleosomal DNA

In the structure of the nucleosome-PU.1 complex, PU.1 recognizes the AAATAGGAA sequence in the canonical motif near the exit site at SHL 5.5 (Fig. 1**d**),^34^ leading to the unwrapping of ∼20 bp nucleosomal DNA. The DNA unwrapping does not lead to the H2A-H2B dissociation (Extended Data Fig. 3**i**). PU.1 interacts with the nucleosomal DNA similarly as with the DNA fragment in the crystal structure (Fig. 2**c**). Notably, in addition to the formation of hydrogen bonds with the GGAA motif, PU.1 ETS residues K171, R222, K245, and K249 also interact with the backbone phosphates beyond the DNA motif and bend the DNA (Fig. 2**c**), bending the DNA towards itself. Similarly, PU.1 also bends the unwrapped nucleosomal DNA (Fig. 2**d**).

The *CX3CR1* DNA also includes GGAA core sequences at SHLs -1.5 and -1.0 in the structure of the nucleosome bound to PU.1 (Fig. 1**a**), which are accessible to PU.1. To investigate the cause for the absence of PU.1 at these two locations in our structures, we conducted EMSA experiments using the free *CX3CR1* DNA and its mutant with the GG<u>AA</u> sequence mutated to GG<u>GG</u> in the two regions. The mutation only slightly reduced the apparent binding affinity of PU.1 (Extended Data Fig. 8). These results explain the absence of PU.1 at these two regions in our structure and reveal that the GGAA motif alone without considering flanking sequences is not a good indicator to predict PU.1 binding although it is the major component of the canonical motif of PU.1.^34^

### Interactions between PU.1 and H2A

In the structural model of the PU.1-nucleosome complex (Fig. 3**a** and Extended Data Fig. 4), PU.1 residue Q218 and H2A residues K75, K76, and T77 in the nucleosome at the interface are close, suggesting that they may form interactions. Notably, PU.1 Q218H mutation is associated with acute myeloid leukemia.^17,18^ To test this hypothesis, we mutated Q218 to His and H2A residues K75, K76 and T77 to Ala (termed ^3A^H2A) and measured the apparent binding affinity changes. We performed EMSA experiments by titrating the nucleosome with PU.1. Fitting the binding data to the Hill equation showed that the mutations of PU.1 Q218H and ^3A^H2A reduced the binding affinity by ∼2- and 3-fold, respectively (Fig. 3**b, c**). In addition, EMSA experiments showed that Q218H has little effect on PU.1 binding to the DNA (Fig. 3**d,e**), further supporting that PU.1 interacts with H2A.

**Fig. 3.**
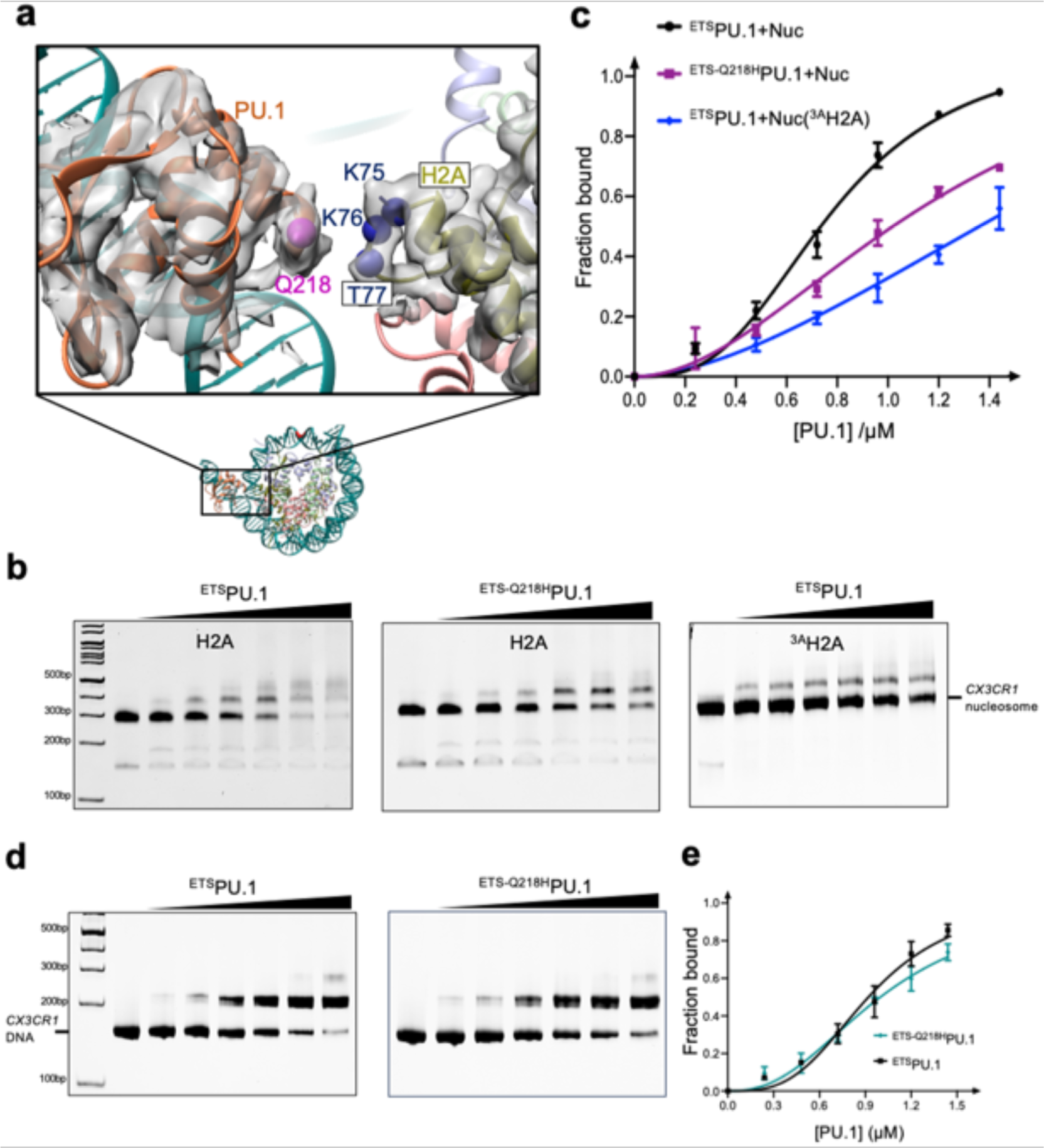
Interactions between PU.1 and H2A. **a**. Density maps for the interface region between PU.1 and H2A in the PU.1-nucleosome complex with an engineered disulfide bond between ^G220C^PU.1 and ^T77C^H2A and the structural model. The residues at the interface are represented by the spheres centered on the C_β_ carbons. **b**. EMSA results of PU1 binding to the nucleosome: ^ETS^PU.1 and the nucleosome containing H2A (left), ^ETS-Q218H^PU.1 and the nucleosome containing H2A (middle), and ^ETS^PU.1 and the nucleosome containing ^3A^H2A. **c**. K_d_^app^ measurement of PU.1 binding to the nucleosome and the effects of mutations using the data from **b**. The lines represent the fitting curves to Hill equation. Data represent mean and s.d. calculated from three independent experiments. **d**. EMSA results of *CX3CR1* DNA binding to ^ETS^PU.1 (left) and ^ETS-Q218H^PU.1 (right). **e**. K_d_^app^ measurement of *CX3CR1* DNA binding to ^ETS^PU.1 and ^ETS-Q218H^PU.1 based on the results in **d**. Data represent mean and s.d. based on the results of three independent experiments.

To investigate the role of these interactions in nucleosome repositioning by PU.1 binding, we performed restriction enzyme digestion using Sau96I. Based on our structures, the cutting site of Sau96I is inside the nucleosome core particle of the free 162 *CX3CR1* nucleosome. However, it will shift to the exposed linker DNA region near the entry site after PU.1 binding to the nucleosome and subsequently increase its accessibility by the restriction enzyme (Fig. 4**a**). Indeed, we found that PU.1 binding increased the cutting efficiency of the 162 bp *CX3CR1* nucleosome (Fig. 4**b**, **c**). We next examined the effects of PU.1 Q218H and ^3A^H2A mutations on digestion efficiency. PU.1 Q218H and ^3A^H2A mutations decreased digestion efficiency compared with wild-type (WT) PU.1 and H2A (Fig. 4**b**, **c**). These results are consistent with the free nucleosome and the nucleosome-PU.1 structures, and reveal that the interactions between PU.1 and H2A play essential roles in the shift of the nucleosomal DNA. To further confirm the DNA repositioning at different temperatures and buffer conditions, we conducted FRET experiments by labeling the H2A residue 116 with Cy3 and the entry nucleotide 1 with Cy5 in the presence of 140 mM KCl or 150 mM NaCl at both 4 and 20 °C (Extended Data Fig. 9). PU.1 binding resulted in a significant decrease in FRET signals under all these conditions.

**Fig 4.**
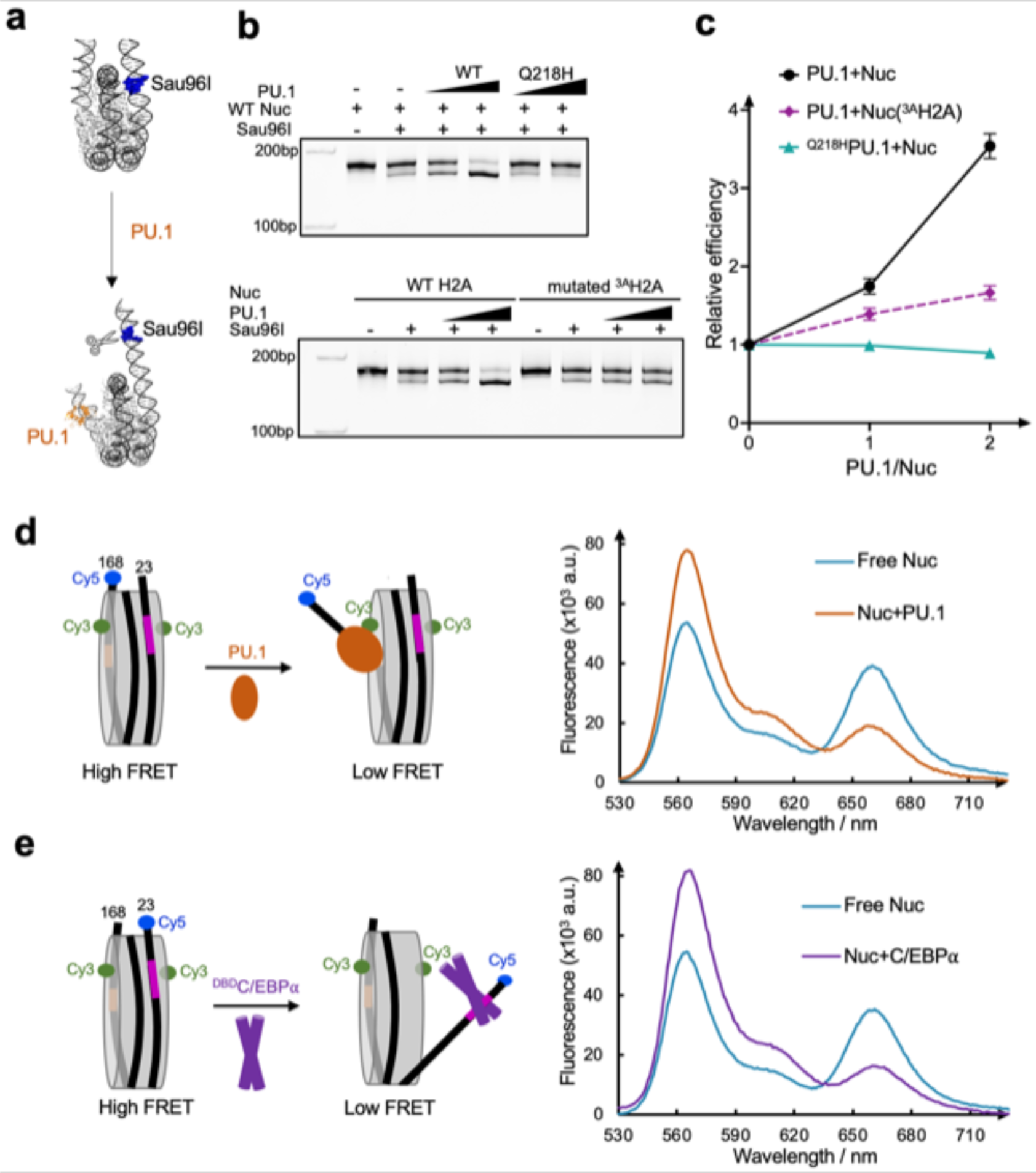
Repositioning and unwrapping of nucleosomal DNA. **a,** Design of restrict enzyme digestion assay. **b**, Restriction enzyme digestion assay of the nucleosome bound to the wild type and mutated PU.1. Each experiment was repeated three times with similar results. **c,** Relative digestion efficiency. Data represent mean and s.d. based on three independent experiments. **d, e.** FRET assays using the 146 bp nucleosome for PU.1 (**d**) and ^DBD^C/EBP**⍺** (**e**). Filled circles (green for Cy3 and blue for Cy5) represent the labeled dyes.

### FRET assay on nucleosomal DNA unwrapping

To validate the unwrapping of the nucleosomal DNA by PU.1 at the exit side, we reconstituted the 146 bp *CX3CR1* nucleosome consisting of nucleotides 23-168. The shorter DNA forces the nucleosome core particle to be in a position mimicking that in the 162 bp nucleosome repositioned by PU.1. Therefore, binding of PU.1 to this 146 bp DNA nucleosome will not cause translation of the DNA, allowing the FRET to report the anticipated DNA unwrapping at the exit site only. We labeled H2A residue 116 with Cy3 and the DNA at the exit site with Cy5. We conducted a FRET assay by titrating PU.1 to the nucleosome (Fig. 4**d**). PU.1 binding decreased the FRET signal, supporting its role in DNA unwrapping at the exit side. We applied a similar assay with the Cy5 labeling at the DNA entry site to investigate the effect of C/EBP⍺ binding on the unwrapping of DNA. We found that the FRET signal decreased with the presence of ^MBP-DBD^C/EBP⍺, confirming that ^MBP-DBD^C/EBP⍺ binding unwrapped the DNA at the entry side (Fig. 4**e**).

### PU.1 facilitates C/EBP⍺ binding to the nucleosome

Our data show that PU.1 binding shifts the C/EBPα binding site from the inner nucleosome core region to the region near the entry site. According to the site-exposure model,^35^ PU.1 binding would facilitate the binding of C/EBP⍺ (Fig. 5**a**). To verify it, we conducted FRET experiments by labeling C/EBP⍺ with Cy5 and the nucleotide 19 with Cy3, which is close to the C/EBP⍺ binding site. Titration of C/EBP⍺ to nucleosome led to a gradual increase of FRET signals, confirming that C/EBP⍺ binds on the unwrapped DNA at the entry side (Fig. 5**b** and Extended Data Fig. 9**e**). Addition of PU.1 led to the increase of FRET signals more rapidly, suggesting that PU.1 facilitates the binding of C/EBP⍺ to the nucleosome, consistent with the earlier *in vivo* result that C/EBP⍺ binding is partially dependent on PU.1.^22,23^ We also conducted a FRET experiment by labeling the DNA at the exit site with Cy3 and PU.1 with Cy5. Titration of PU.1 to nucleosome increased the FRET signal (Fig 5**c** and Extended Data Fig. 9**f**). However, the addition of C/EBPα did not facilitate PU.1 binding to the nucleosome.

**Fig 5.**
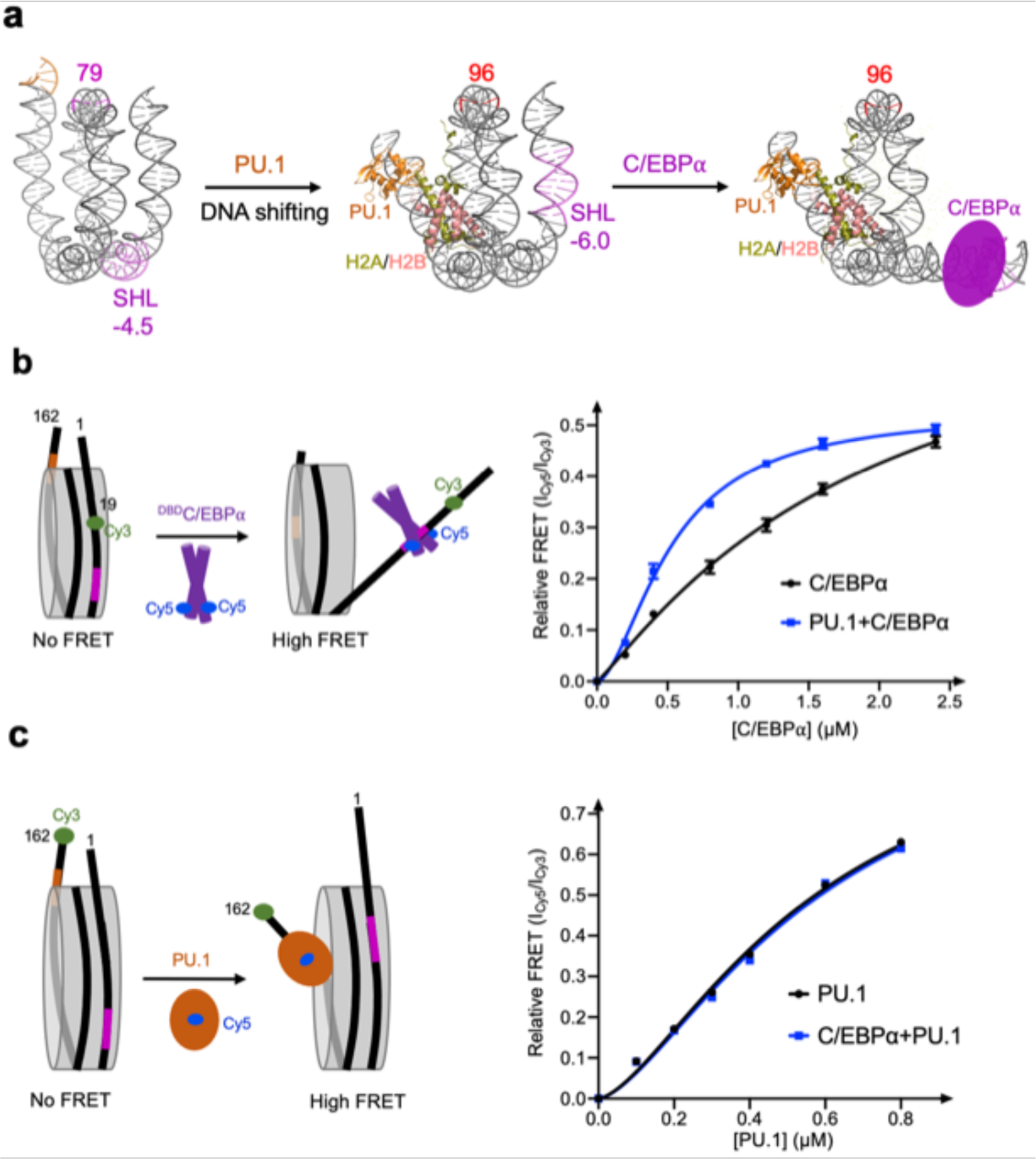
PU.1 facilitates C/EBPα binding to nucleosome. **a.** Diagram shows the process of subsequent binding of two pioneer factors. **b.** Binding of C/EBPα to nucleosome without and with PU.1. The relative FRET intensity is calculated by the ratio of Cy5 peak intensity at 665nm and Cy3 peak intensity at 565nm. **c.** Binding of PU.1 to nucleosome without and with C/EBPα. The data in **b** and **c** represent mean and s.d. based on three independent experiments.

### H1 eviction by PU.1 and C/EBP⍺

The unwrapping of nucleosomal DNA by PU.1 and C/EBP⍺ at both exit and entry sides suggest that the two pioneer factors together might evict linker histone in the chromatosome (Fig. 6**a**). We tested this hypothesis using FRET by labeling Cy3 on H2A and Cy5 on ^K26C^H1.4. Neither PU.1 nor ^MBP-DBD^C/EBP⍺ caused a significant change in the FRET signal. In contrast, together they led to a large decrease in the FRET signal (Fig. 6**b**), indicating that the synergistic binding of PU.1 and C/EBP⍺ unwraps nucleosomal DNA from both sides of the nucleosome, causing H1 eviction. Moreover, PU.1 Q218H mutant showed a much weaker H1-eviction function, consistent with our earlier results that the interactions between PU.1 and H2A play essential roles in PU.1 binding and nucleosome repositioning (Fig. 6**c**).

**Fig 6.**
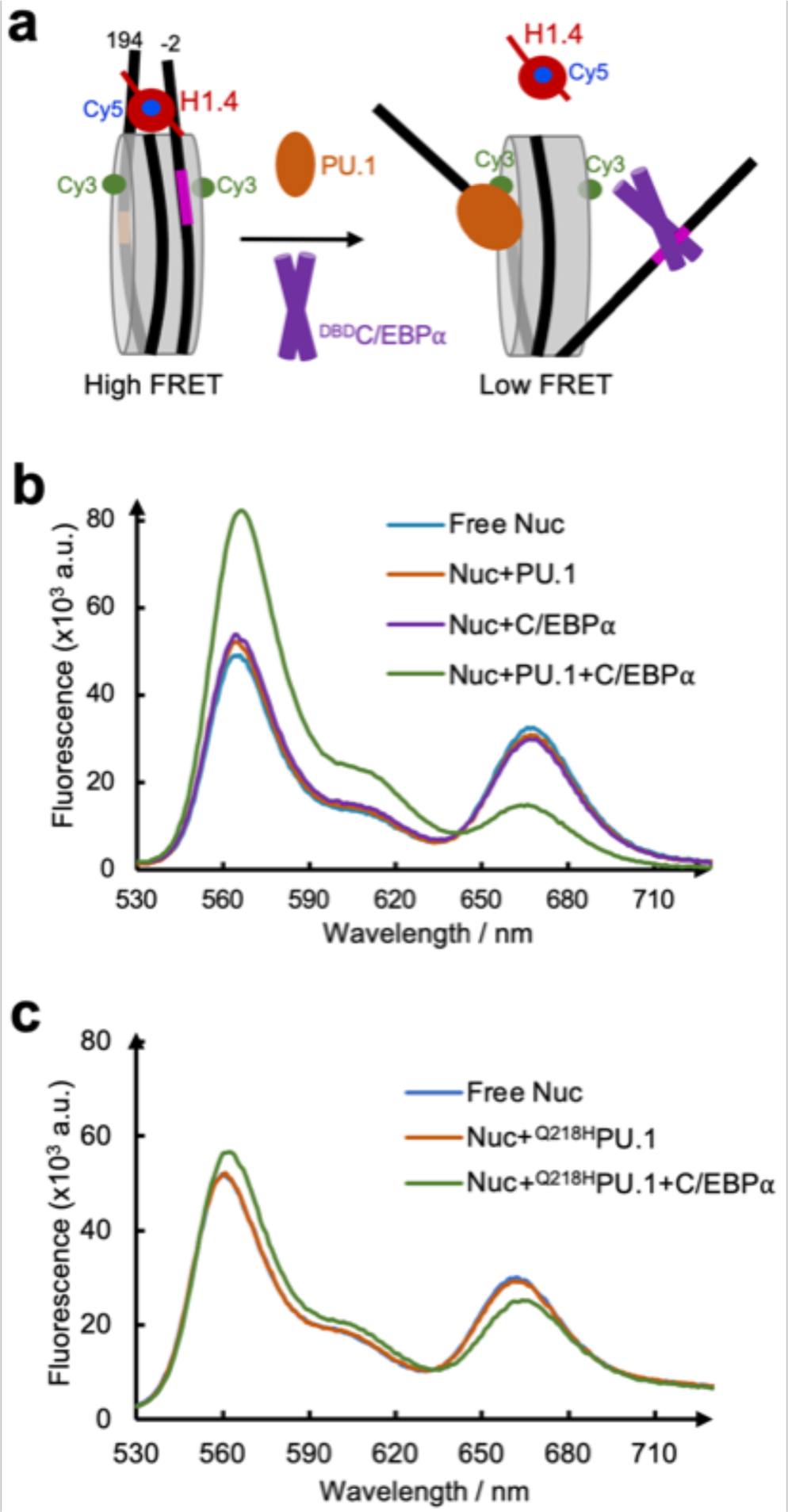
FRET assay on the eviction of H1 by PU.1 and ^DBD^C/EBP⍺. **a**. FRET assay design. **b.** FRET assay showing the effect of PU.1, ^DBD^C/EBP⍺ and a combination of two pioneer factors on H1 eviction. **c.** FRET assay showing the effect of PU.1 mutant Q218H on H1 eviction.

### Opening of H1-condensed nucleosome array by PU.1 and C/EBP⍺

Pioneer transcription factors can open closed chromatin without using ATP. To verify it, we engineered XhoI and EcoRI cutting sites near the binding sites of PU.1 and C/EBP⍺, respectively, in the *CX3CR1* nucleosome (Fig. 7**a**, Extended Data Fig. 10**a**). We then reconstituted a 12 x 197 bp nucleosome array (NA) with five and six W601 nucleosomes on each side of the engineered *CX3CR1* nucleosome (Fig. 7**b**, Extended Data Fig. 10). We condensed the nucleosome array using H1. Finally, we performed the restriction enzyme digestion experiments with an increasing amount of PU.1 or C/EBP⍺ or both. Under the same experimental conditions, we found that PU.1 and C/EBP⍺ facilitate the restriction enzymes to cut the nucleosome array (Fig. 7**c, d**), suggesting they can open the closed chromatin. In contrast, the nucleosome array with the *CX3CR1* DNA replaced with W601 showed resistance to the restriction enzymes, indicating that the opening only occurred locally at the *CX3CR1* nucleosome (Extended Data Fig. 10**f, g**). The enzyme cutting efficiency did not increase using the nucleosome array with mutations at PU.1 binding site and C/EBP⍺ binding site (Fig 7**c,d**, Extended Data Fig. 10**h, i**), suggesting that both binding sites are indispensable for the nucleosome array opening. Compared with C/EBP⍺, PU.1 opened the nucleosome array more efficiently. Moreover, we found that PU.1 can facilitate C/EBP⍺ to open the nucleosome array (Fig. 7**e**), in agreement with the FRET result (Fig. 5**b**).

**Fig 7.**
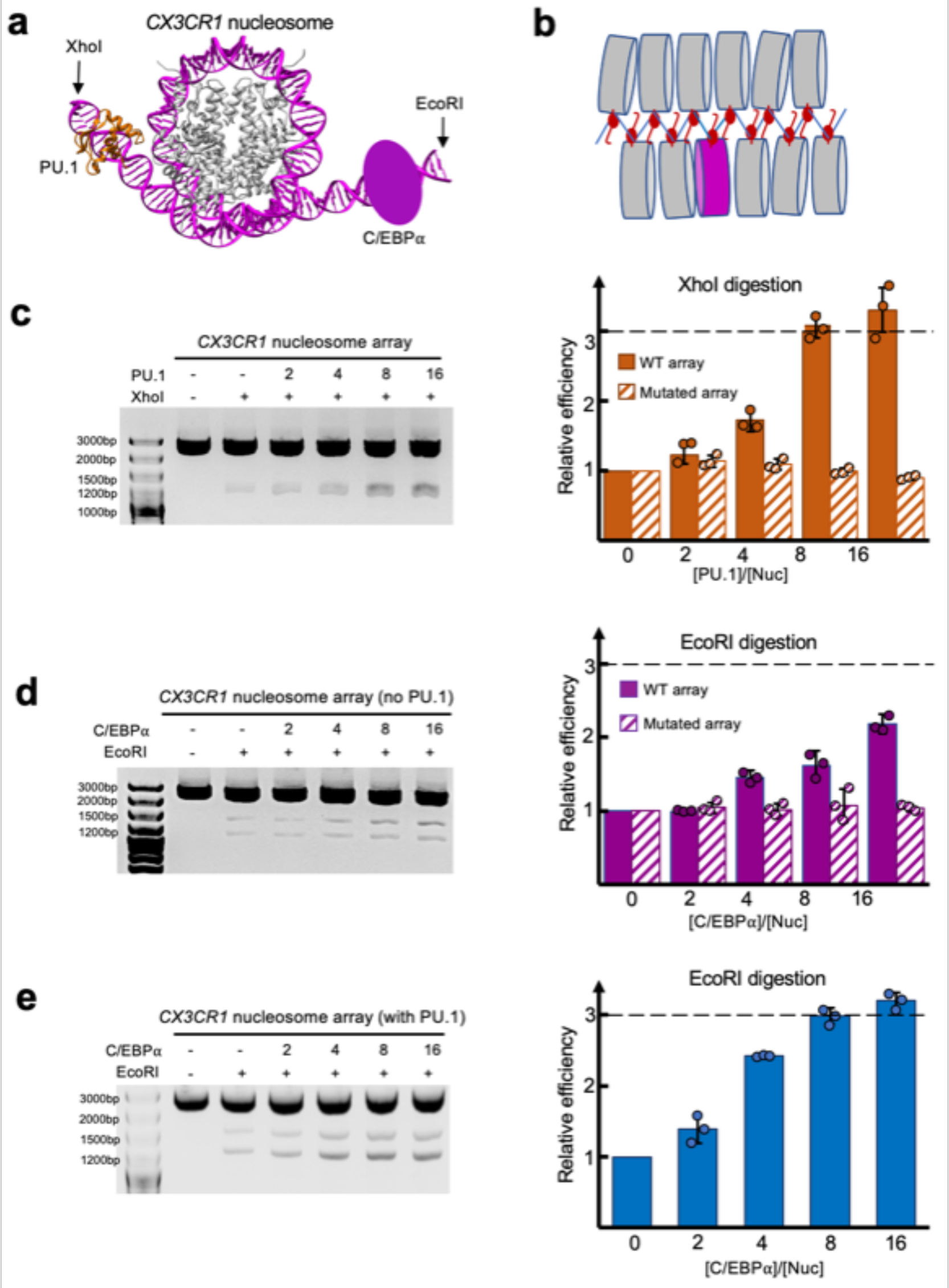
Opening of H1-condensed nucleosome array by PU.1 and C/EBP⍺. **a.** Illustration of engineered EcoRI and XhoI cutting sites in the *CX3CR1* DNA based on the nucleosome-PU.1-^DBD^C/EBP⍺ structural model, which is proximal to C/EBP⍺ and PU.1 binding site, respectively. **b.** Illustration of the 12 x 197 bp nucleosome array with the *CX3CR1* nucleosome in the middle among the ‘W601’ nucleosomes. **c.** Digestion efficiencies of the nucleosome arrays with WT (solid bar) and mutated DNA (GGAA to GGGG at PU.1-binding site) (bar with slashed lines) by XhoI at multiple PU.1 concentrations. **d**. Digestion efficiency of the nucleosome arrays with WT (solid bar) and mutated DNA (CAGC<u>TGG</u>TTG to CAGC<u>AAC</u>TTG at the C/EBP⍺-binding site) (bar with slashed lines) by EcoRI at multiple C/EBPα concentrations. **e.** Digestion efficiency of the nucleosome array by EcoRI at multiple C/EBPα concentrations in the presence of PU.1 (the ratio of PU.1 over the nucleosome array is 4). In **c**-**e**, data represent mean and s.d. based on the results of three independent experiments.

## Discussion

Nucleosomes were thought inhibitory for binding transcription factors due to potential steric obstructions from core histones and the neighboring DNA gyres. To explain how transcription factors may bind the nucleosome, the site-exposure model hypothesizes that nucleosomal DNA may spontaneously unwrap and expose the DNA motifs for transcription factor binding.^35^ On the other hand, functional genomics and structural modeling studies suggest that the pioneer transcription factor functionality may be determined by how they can avoid the steric clash with the nucleosome when recognizing the exposed DNA motif.^36^ Recent structural studies using nucleosomes with non-genomic DNA as models suggest that the DBDs of transcription factors can bind the exposed DNA motifs in the nucleosome core region with or without DNA unwrapping.^10,11^

Our results show that PU.1 and C/EBP⍺ each can bind the *CX3CR1* nucleosome and unwrap nucleosomal DNA, suggesting that they may use the site exposure mechanism for targeting the nucleosome *in vivo*. In addition, the interactions between PU.1 ETS and H2A loop indicate that PU.1 may stabilize the PU.1-nucleosome complex and possibly decrease its dissociation rate,^8,37^ which would increase the residence time of pioneer factors on the nucleosome. For C/EBP⍺ binding at the entry site, structural modeling suggests that C/EBP⍺ may only bind the unwrapped DNA as its binding to fully wrapped nucleosome would cause steric clashes with the neighboring DNA gyre and core histones (Extended Data Fig. 6**i**). Notably, EMSA experiments revealed that when the ratio of pioneer factor to nucleosome is high, multiple PU.1 and C/EBP⍺ molecules can bind to the CX3CR1 nucleosome (as shown in Extended Data Figs. 1**b, d**). These additional bindings are likely the result of C/EBP⍺ associating with the nucleosome at weaker binding sites. The investigation of these higher-order bindings and the relevance of our results to the *in vivo* function using genome editing is a topic for future studies. However, it is important to note that these findings should not impact the roles of PU.1 and C/EBP⍺ as identified in our structures. This is because the mutations observed in the DNA sites bound to the pioneer factors in our structures can effectively eliminate the ability of the pioneer factors to open the H1-condensed nucleosome arrays (as depicted in Fig. 7**c**, **d**).

Our study reveals a mechanism for the cooperative binding of two pioneer factors whereby one pioneer factor (PU.1) binds the DNA motif and core histones, leading to the repositioning of the nucleosome, which in turn shifts the DNA motif of the other transcription factor (C/EBP⍺) from the inner nucleosome core region to the location near the entry site (Fig. 5 and Supplementary Data Movie 1). Our findings indicate that the DNA within the CX3CR1 nucleosome exhibits relatively weaker interactions with the core histones. In vivo, the movement of the nucleosomal DNA may be further facilitated by chromatin remodelers. Recent studies have demonstrated that the intrinsically disordered region of PU.1 can recruit chromatin remodelers, promoting the opening of chromatin.^38^ Additionally, the binding between PU.1 and H2A can provide stabilizing energy to reposition the nucleosome, which can be enhanced through mass actions with increased concentrations of transcription factors. It should be noted that the local concentrations of pioneer factors at enhancer regions remain largely unknown, but they could potentially be dramatically increased through the formation of condensates. For instance, OCT4 and Med1 interact through liquid-liquid phase separation to form condensates,^39^ significantly elevating the local concentration of OCT4 at enhancer sites. Moreover, recognizing the nucleosome likely necessitates high local concentrations of pioneer factors, given that the concentration of the CX3CR1 nucleosome in the cell is expected to be extremely low.

Previous studies using restriction enzymes and DNA unzipping have shown that the location near the entry/exit site has a higher probability of unwrapping spontaneously than in the inner region of the nucleosome core particle.^40,41,35^ Thus, PU.1 appears to have taken advantage of this physical mechanism to facilitate the binding of C/EBP⍺ by repositioning the nucleosome for their pioneer functions. Furthermore, this mechanism could explain the *in vivo* results that PU.1 alone can target compacted chromatin and induce its accessibility, whereas C/EBP⍺ binding to nucleosome-enriched regions partially depends on PU.1.^22,23^ Finally, the synergistic binding mechanism for the two transcription factors revealed from our study is distinct from known models for cooperative binding of transcription factors to DNA,^42,43^ including direct interactions between transcription factors, DNA-mediated interactions, and nucleosome displacement.

## Acknowledgments

We thank Dr. Rick Huang and Ms. Allison Zeher for assistance in cryo-EM Data collection. The cryo-EM work utilized NCI-NIH IRP Cryo-EM Consortium (NICE) microscopy resource and NIH high performance computing Biowulf system for data processing. This work is supported by the intramural research program at the Center for Cancer Research, National Cancer Institute, National Institutes of Health (Y.B.).

## Author contributions

Y.B. conceived the project. T.L. and R.G. conducted the experiment. B-R. Z. provided the scFv, H1 protein, and W601 array DNA. Y.B., R.G., and T.L. analyzed the structures and wrote the paper.

## Methods

### Expression and purification of histones

Recombinant human histones and their mutants were expressed individually in *Escherichia coli* BL21(DE3) cells. All mutations were generated using QuikChange kit (Agilent). E. coli cells harboring each histone expression plasmid were grown at 37 °C in 2 x YTB Broth. When OD_600_ reached around 0.6-0.8, 0.5 mM IPTG was added to induce recombinant protein expression for 5 h at 37 °C. The cells were harvested and resuspended in 50 ml of buffer A (50 mM Tris-HCl, 500 mM NaCl, 1 mM PMSF, 5% glycerol, pH 8.0), followed by sonication on ice for 360s with a pulse of 15s on and 30s off. The cell lysates were centrifuged at 40,000 x g for 20 min at 4 °C. The pellet containing histones was resuspended in 50 ml of buffer B (20 mM sodium acetate, pH 5.2, 100 mM NaCl, 5 mM 2-mercaptoethanol, 1 mM EDTA, and 6 M urea). The samples were rotated for 12 h at 4 °C, and the supernatant was recovered by centrifugation at 96,000 x g for 60 min at 4 °C. The supernatant was loaded to Hitrap S column chromatography (GE Healthcare). The column was washed with buffer B, and histone protein was eluted by a linear gradient of 100 to 800 mM NaCl in buffer B. The purified histones were dialyzed against water three times and freeze-dried.

### Expression and purification of pioneer transcription factors

The plasmids (pET28a) containing the mouse full-length PU.1 and PU.1 ETS (^ETS^PU.1, residue 165-272) with N-terminal his tag and DBD of C/EBP⍺ (residues 214-359) with C-terminal His-tag were directly synthesized in Genscript. Q218H and G220C mutations were introduced into both full-length PU.1 and ^ETS^PU.1 using QuikChange kit (Extended Data Table 2). The plasmids were transformed into BL21(DE3) cells and grown in LB medium. When OD_600_ reached 0.8, 0.5 mM IPTG was added at 37 °C for 2 hrs. Expression and purification procedures was finished within 1 day for the full-length protein to avoid degradation. Cells were harvested and resuspended in the buffer containing 20 mM HEPES, pH 7.1, 1 M NaCl, 0.5 mM TCEP, and a protease inhibitor cocktail (Roche). After sonification and ultracentrifuge, the supernatant went through the column containing nickel affinity beads (Thermal Fisher). The column was washed with the buffer containing 20 mM HEPES, pH 7.1, 1 M NaCl, 0.5 mM TCEP, 80 mM imidazole, then eluted by the buffer containing 20 mM HEPES, pH 7.1, 1 M NaCl, 0.5 mM TCEP, 250 mM imidazole. The solution was concentrated to ∼1 ml and went through gel filtration on Superdex 75 10/300 increase column (GE Healthcare) equilibrated with a buffer containing 20 mM HEPES, pH 7.1, 200 mM NaCl, 0.5 mM TCEP. C/EBP⍺ DNA binding domain containing residues 214-359 (^DBD^C/EBP⍺) was coned both into pET28a with C-terminal His-tag and pET30a with an N-terminal MBP tag (^MBP-DBD^C/EBP⍺). I295C/C357A was introduced into ^DBD^C/EBP⍺ for the labeling of fluorescence dye. All C/EBP⍺ proteins were expressed and purified the same way as PU.1.

### Preparation of DNA

DNA fragments (Supplementary Data Table 1) were prepared by PCR amplification, followed by ethanol precipitation and purification using the POROS column. Briefly, The PCR products were pelleted by 75% ethanol containing 0.3 M NaAc at pH 5.2. The sample was incubated for 120 min at -20 °C, followed by centrifugation. The pellet was resuspended by TE buffer, loaded to POROS column chromatography (GE Healthcare), and washed with buffer containing 20 mM Tris-HCl, pH 7.4, 5 mM 2-mercaptoethanol. The DNA was eluted by a linear gradient of 0 to 2 M NaCl and stored at 4 °C.

### Reconstitution of nucleosomes and nucleosome arrays

Purified recombinant histones in the equal stoichiometric ratio were dissolved in ∼6 ml unfolding buffer (7 M guanidine-HCl, 20 mM Tris-HCl, pH 7.4, 10 mM DTT) and were dialyzed against refolding buffer (10 mM Tris-HCl, pH 7.4, 1 mM EDTA, 1 mM DTT, 2 M NaCl) for about 12 hours. The mixture was centrifuged at 20,000 x g to remove any insoluble material. Soluble octamers were purified by size fractionation on a Superdex 200 10/300 increase gel filtration column. Purified histone octamers and DNA were mixed with a 1.3:1 ratio in high-salt buffer (2 M NaCl, 10 mM K/Na-phosphate, pH 7.4, 1 mM EDTA, 5 mM DTT). 1 ml mixture in a dialysis bag was placed in 600 mL of the high-salt buffer and dialyzed for 30 min followed by salt gradient dialysis. 3 L of a low-salt buffer (150 mM NaCl, 10 mM K/Na-phosphate, pH 7.4, 1 mM EDTA, 2 mM DTT) were gradually pumped into dialysis buffer with a flow rate of 2 ml/min for 18 h. It was then dialyzed against a low-salt buffer for 30 min in the cold room. The sample was then incubated at 37 °C for 3-5 h. The mixture was centrifuged at 20,000 x g to remove any insoluble material. The nucleosomes were further purified by ion-exchange chromatography (TSKgel DEAE, TOSOH Bioscience, Japan) to remove free DNA and histones. The purified nucleosomes were dialyzed against the buffer containing 20 mM HEPES, pH 7.1, 0.5 mM TCEP.

To reconstitute a 12x197 bp nucleosome array with the *CX3CR1* nucleosome in the middle, a pUC-18 plasmid was reconstructed with the incorporation of sequentially ligated fragments of 5 x 197 bp W601 DNA, 1 x 197bp *CX3CR1* DNA, and 6 x 197 bp W601 DNA (Extended Data Fig. 10). The resulting 12 x197 bp DNA in the plasmid was verified by Sanger sequencing and restriction digestion. Large-scale preparation and purification of the plasmid followed the previous protocol.^44^ Briefly, 12 x 197 bp DNA was released from the plasmid by EcoRV digestion (150 unit enzyme per 1 mg plasmid) and purified by stepwise PEG 6000 precipitation. The fractions containing the 12 x 197 bp DNA was used for nucleosome array reconstitution according to early methods.^45^ Saturation of nucleosome in the reconstituted nucleosome array was verified by SmaI digestion, which showed a predominant mono-nucleosome band on a 1.2% agarose gel. Linker histone H1.4-bound nucleosome array was made by mixing 1.3-fold of H1.4 (relative to the molar concentration of nucleosomes in the array) with the nucleosome array in 10 mM Tris-HCl, pH 8.0, 1 mM EDTA, 1 mM DTT and 0.6 M NaCl buffer, followed by dialysis using the same buffer without NaCl.

### Cryo-EM sample preparation and data collection

2 μM (50 μL) nucleosome containing mouse *CX3CR1* DNA (162 bp), 6 μM (50 μL) ScFv, and 40 μM (20 μL) PU.1 were mixed in the buffer of 20 mM HEPES, pH 7.1, 150 mM NaCl, 0.5 mM TCEP. The sample was incubated on ice for 0.5 h and then concentrated to 200 ng/ul (measured by OD_260_), which was loaded onto the glow-discharged holey carbon grid (Quantifoil 300 mesh Cu R1.2/1.3). For the disulfide bond mutation complex, nucleosomes containing ^T77C^H2A and ^G220C^PU.1 were used at the same concentration instead of wild-type nucleosomes and wild-type PU.1. DNA was elongated to 167 bp with 5 additional base pairs ‘TTCTA’ at the exit site for better PU.1 binding, and 2 mM oxidized glutathione was added to the complex solution to accelerate disulfide bond formation. For nucleosome-ScFv-PU.1-C/EBP⍺ quaternary complex, 2 μM (50 μL) nucleosome containing mouse *CX3CR1* DNA (167 bp) and ^T77C^H2A, 6 μM (50 μL) ScFv, and 40 μM (20 μL) ^G220C^PU.1 and 40 μM (40 μL) dimeric ^MBP-DBD^C/EBP⍺ were mixed in the buffer of 20 mM HEPES, pH 7.1, 200 mM NaCl, 0.5 mM TCEP. The sample was then concentrated to 100 ng/ul (measured by OD_260_). Holey Lacey carbon grids (Tedpella, 300 mesh, Cu) were used. The grids were blotted for 3 s at 14 °C and 100% relative humidity using an FEI Vitrobot Mark IV plunger before being plunge-frozen. For the free 162bp *CX3CR1* nucleosome sample, an equal volume (25 μL) for each 4 μM nucleosome was mixed with 12 μM scFv and the grids were prepared using the same parameter for the nucleosome-PU.1 complex.

Cryo-EM data were collected using a Titan Krios G3 electron microscope (Thermo-Fisher) operated at 300kV. Micrographs were acquired in super-resolution mode at the nominal magnification of 81,000x with 0.528 Å image pixel size using a 20-eV slit post-GIF Gatan K3 camera. The dose rate on the camera was set to 15 e^-^/pixel/s. The total exposure time of each micrograph was 4 s fractionated into 50 frames with 0.08 s exposure time for each frame. The data collection was automated using the SerialEM software package.^46^ A total of 2,986 micrographs were collected for free *CX3CR1* nucleosome sample, and 4,503 micrographs were collected for the wild-type nucleosome-PU.1 complex sample. 7,014 micrographs were collected for the disulfide bond mutated nucleosome-^G220C^PU.1 sample and 5,204 micrographs were collected for the nucleosome-^G220C^PU.1-^MBP-DBD^C/EBP⍺ complex sample.

All datasets were processed in RELION/3.1.3 and cryoSPARC v3.2 following the standard procedures.^47,48^ The beam-induced image drift was corrected using MotionCor2.^49^ The averaged images without dose weighting were used for defocus determination using CTFFIND4.1^50^, and images with dose weighting were used for particle picking and extraction. Particles were automatically picked using Gautomatch (https://www.mrc-lmb.cam.ac.uk/kzhang/Gautomatch/).

1,274,643 particles were picked for the free *CX3CR1* nucleosome dataset. Bad particles were removed by 2D classification and 3D classification in RELION using 4x binned particles. 674,406 particles were selected and re-extracted without binning, followed by two more rounds of 3D classification using 7.5° and 3.7° angular sampling rate. 1 class containing 143,068 particles with good density of DNA was selected for auto-refine. Bayesian Polishing, followed by importing to CryoSPARC for CTF-refinement and non-uniform refinement, further improved the resolution to 2.6 Å.

For Nuc-ScFv-PU.1 complex dataset, 2,506,660 particles were picked from 4,503 micrographs. Bad particles and free nucleosome particles were removed from 2D classification and 3D classification using 2x binned particles. 581,016 particles with one blurry DNA end were selected and re-extracted without binning. Two more rounds of 3D classification with angular sampling rate 7.5° and 3.7° were performed. one class with good density of unwrapped DNA end was selected for focused 3D classification without alignment and with a small mask only covering the unwrapped DNA part. Another two classes with blurry DNA end density were also selected for focused 3D classification without alignment. The two focused 3D classification jobs generate two classes with clearly unwrapped DNA and also the extra bulb density on the unwrapped DNA. The two classes were combined and submitted for Bayesian Polishing. After CTF-refinement and 3D-auto refinement in RELION, a 2.9 Å map was generated for model building. The PU.1 ETS domain (PDB: 1PUE) can fit into the extra density on the unwrapped DNA well.

For the complex containing nucleosome (^T77C^H2A), scFv, ^G220C^PU.1 complex, 3,243,708 particles were picked and submitted for 2D classification using 4x binned particles. 2,000,352 particles were selected for 3D classification using the above wild type nucleosome-scFv-PU.1 map lowpass filtered into 60 Å as the initial model. 622,795 particles with both good nucleosome density (contained one side unwrapped DNA) and also good ^ETS^PU.1 density were selected. Particles were then re-extracted without binning and submitted to 2 more rounds of 3D classification. One class with high-resolution feature and good ^ETS^PU.1 density were selected. Bayesian Polishing was performed, followed by importing into CryoSPARC. CTF refinement and non-uniform refinement were used to further improve the resolution to 2.7 Å.

For the nucleosome (^T77C^H2A)-scFv-^G220C^PU.1-^MBP-DBD^C/EBP⍺ complex dataset, 2,888,287 particles were picked, and 1,411,125 particles were selected after 2D classification using 4x binned particles. 3D classification using the above nucleosome (^T77C^H2A)-scFv-^G220C^PU.1 lowpass filtered to 60 Å as the initial model generated a class with good nucleosome and PU.1 density. 468,250 particles in this class were then re-extracted without binning, and two more rounds of 3D classification were performed. 1 class with good PU.1 density and clearly unwrapped entry site DNA density was selected for 3D auto-refine, generating a 3.9 Å map with 116,597 particles. Focused refinement with mask only covering the nucleosome core and PU.1, excluding the unwrapped entry site DNA, generated a 2.8 Å map. The focused 3D classification without alignment and with mask only covering the unwrapped entry site DNA generated three classes all around 4.1 Å. For these 3 classes, the entry site DNA was all unwrapped.

### Model building and structure analysis

For the free *CX3CR1* nucleosome, an initial model of the nucleosome histone octamer and scFv was generated using the nucleosome structure (PDB: 7K61). The model was fitted into the cryo-EM density map of the *CX3CR1* nucleosome-scFv complex. The DNA sequence was built into the map from scratch in COOT ^51^ and the histone octamer and scFv were optimized by manual rebuilding. The whole complex was refined using real-space refinement in PHENIX.^52^ For the nucleosome-scFv-PU.1 complex, the free *CX3CR1* nucleosome-scFv was used as the initial model. Histone octamer and scFv were fitted into the map and manually rebuilt in COOT, and DNA was built in COOT. The model was refined using real-space refinement in PHENIX. ^ETS^PU.1 from the crystal structure (PDB: 1PUE) was rigid-body fitted into the density on the unwrapped DNA. The side chains including I217 and Q218 were adjusted based on the interaction with H2A. For the nucleosome (^T77C^H2A)-scFv-^G220C^PU.1 complex, the nucleosome-scFv-PU.1 was used as the initial model. The whole model was rigid-body fitted into the map, and both proteins and DNA were manual rebuilt in COOT. The model was further optimized using real space refinement in PHENIX. For the nucleosome (^T77C^H2A)-scFv-^G220C^PU.1-^MBP-DBD^C/EBP⍺ complex, the nucleosome-scFv-PU.1 structure was used as the initial model. Histone octamer, PU.1 and scFv were fitted into the map, and DNA was built in COOT. The structure was optimized by manually rebuilding in COOT, followed by further refinement using real-space refinement in PHENIX. C/EBP⍺ model from the structure (PDB 1NWQ) was rigid-body fitted into the extra density on the unwrapped entry site DNA. All the figures and movies were made using UCSF Chimera^53^ and PyMOL (Version 1.8, Schrödinger, LLC. DeLano Scientific).

### Electrophoretic mobility shift assay

Nucleosomes or free DNA at a final concentration of 0.5 μM were mixed with purified proteins at different concentrations. Typical binding reactions were carried out for 30 min at room temperature in buffer containing 20 mM HEPES, pH 7.3, 150 mM NaCl, and 0.5 mM TCEP. Six μl of the binding reactions were analyzed on 4% or 5 % acrylamide gels in 0.2 x TBE at 120 V for 60 minutes at 4 °C. After electrophoresis, gels were stained with ethidium bromide (EtBr) and quantified using ImageJ. For the apparent binding affinity measurements, we quantified band intensities using ImageJ, subtracted against the background and normalized to the signal of the input free nucleosome. We used Prism (GraphPad) to fit the binding data with the Hill equation: ε = [L]^n^/(K_d_^app^ + [L]^n^). Here, ε is the fraction of total nucleosome bound to PU.1 or C/EBPα (containing all the shifted bands). [L] is the total concentration of corresponding pioneer factors. K_d_^app^ is the apparent binding constant. n is the Hill coefficient.

### DNA repositioning experiments

180bp native *CX3CR1* nucleosome (Table S2) was used in this assay. 1000 ng nucleosome was mixed with PU.1 at different concentrations in 18.5 μl reaction system in buffer 20 mM HEPES, pH 7.1, 100 mM NaCl and 0.5 mM TCEP. After 20 min incubation at room temperature, 1 μl Cutsmart buffer and 1 U Sau96I (New England Biolabs) were added into the system. The mixture was placed into 37 °C incubator for 20 min. Purple gel loading dye (New England Biolabs) was added to stop the enzyme digestion reaction. Then 1 μl proteinase K (New England Biolabs) was used at 50 °C for 1 hour to digest PU.1 and histones. 4 μl of the digestion products were analyzed on 5 % acrylamide gels in 0.2 x TBE at 120 V for 60 minutes at 4 °C. After electrophoresis, gels were stained with ethidium bromide (EtBr) and quantified using ImageJ. Three independent experiments were performed. Microsoft Excel and Prism were used to generate the figures.

### Fluorescence resonance energy transfer (FRET) assay

^G220C^PU.1, ^MBP-DBD^C/EBP⍺ containing I295C and ^K26C^H1.4 was labeled with Cy5 following the manufacture’s protocol (Cytiva). 1 mg target protein was dissolved in 1 mL degassed buffer containing 20 mM HEPES, pH 7.1, 400 mM NaCl and 1 mM TCEP. One vial of Cy5 maleimide was dissolved in dimethylformamide (DMF) and mixed with target protein. The mixture was agitated overnight at cold room. Size-exclusive chromatography was used to separate the free Cy5 dye. ^L116C^H2A was labeled with Cy3 according to the previous publication.^54^ 1 mg ^L116C^H2A was dissolved in 1 mL degassed buffer containing 800 mM HEPES, pH 7.1, 1.5 mM Guanidine HCl, 1 mM TCEP. 1vial Cy3 maleimide (Cytiva, California) in DMF was mixed with ^L116C^H2A and incubated at room temperature for 5 hours. A Sephadex G-25 column was used to separate the free Cy5 dye. Labeled ^L116C^H2A was refolded to octamer following the same protocol with unlabeled histones.

Fluorescent DNA fragments were produced using PCR by the Cy3 or Cy5 labeled primers (IDT). For the unwrapping assay, PU.1 or ^MBP-DBD^C/EBP⍺ was titrated into 200 nM nucleosome in buffer 20 mM HEPES, pH 7.1, 100mM NaCl, 0.5mM TCEP. For the DNA repositioning assay, PU.1 was titrated into 200 nM nucleosome in buffer 20 mM HEPES, pH 7.1, 150mM NaCl or 20 mM HEPES, pH 7.1, 140mM KCl at both room temperature (20 ℃) and low temperature (4 ℃). For the binding of C/EBP⍺ to 162bp CX3CR1 nucleosome, Cy5 labeled C/EBP⍺ was titrated into 200 nM Cy3 labeled nucleosome (Cy3 on nucleotide 19) in buffer 20 mM HEPES, pH 7.1, 100mM NaCl, 0.5mM TCEP without or with 800 nM unlabeled PU.1. For the binding of PU.1 to 162bp nucleosome, Cy5 labelled PU.1 was titrated into 200 nM Cy3 labeled nucleosome (Cy3 on nucleotide 162) without or with 2 μM C/EBP⍺. In H1 eviction FRET assay, Cy5 labeled ^K26C^H1.4 was mixed with nucleosome containing Cy3 labeled H2A at a ratio of 1.3:1 before adding PU.1, ^Q218H^PU.1 and ^MBP-DBD^C/EBP⍺. The fluorescence intensity was recorded using QuantaMaster from Photon Technology International. The excitation wavelength was set to 510 nm, and emission spectra were collected between 530nm ∼ 730nm. Three independent experiments were performed.

### Restrict enzyme digestion of H1.4-condensed nucleosome array

3 mg wild-type CX3CR1 nucleosome array or mutated nucleosome array or 601 nucleosome array in 20 μL was mixed with PU.1 or ^DBD^C/EBP⍺ with a molar ratio to the nucleosome array ranging from 0:1 to 16:1, respectively, in digestion buffer (10 ul, 10 mM Tris-HCl, pH 8.0, 60 mM NaCl, 1 mM magnesium chloride, 2 mM DTT) at room temperature. 10 units of restriction enzyme XhoI (NEB) were added to the nucleosome array with and without PU.1. 5 units of restriction enzyme EcoRI (NEB) were added to the nucleosome array in the absence and presence of ^DBD^C/EBP⍺. Samples were incubated at 37 °C for 30 min, and the enzyme was inactivated by NEB purple loading dye. Samples were then incubated with proteinase K at 50 °C for 60 min. After centrifugation, the top solution was harvested and loaded into a 1 % agarose gel stained with SYBR Safe dye (Invitrogen). Electrophoresis was performed at 130 V in 1 x TBE buffer for 25 min. Band intensities for the digestion product and input were measured using ImageJ. The relative digestion efficiency was calculated using the intensity of the product in the presence of transcription factors divided by the product intensity without transcription factors.

## Data availability

The cryo-EM reconstructions and atomic models of the *CX3CR1* nucleosome and the nucleosome-pioneer factors complexes have been deposited in the Electron Microscopy Data Bank and the Protein Data Bank under the following accession codes: EMD-40889 and PDB ID 8SYP for the 162 bp *CX3CR1* DNA nucleosome; EMD-28629 and PDB ID 8EVH for the wild type nucleosome-PU.1 complex; EMD-28630 and PDB ID 8EVI for the nucleosome containing ^T77C^H2A bound ^G220C^PU.1; EMD-28631 and PDB ID 8EVJ for the nucleosome (^T77C^H2A) bound ^G220C^PU.1 and ^MBP-DBDR^C/EBPα. Structures used for model building can be found in the PDB, including PDB ID 7K61 for nucleosome bound to scFv, PDB ID 1PUE for PU.1 bound to DNA and PDB ID 1NWQ for C/EBPα bound to DNA.

---

**Extended Data Fig. 1.**
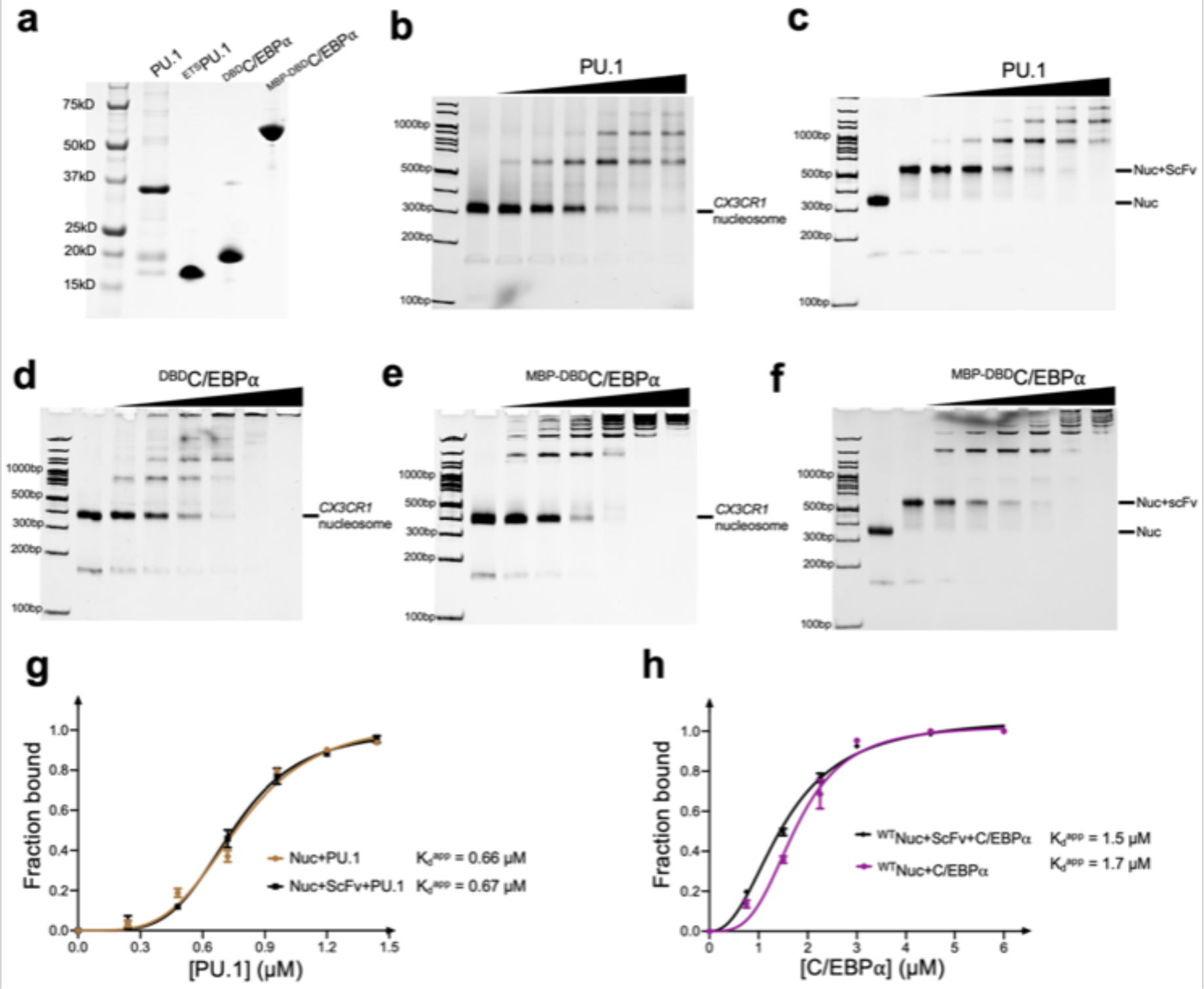
| Characterization of PU.1 and C/EBPα binding to *CX3CR1* nucleosome. **a,** SDS-PAGE of the protein samples used in this study. **b and c,** EMSA of *CX3CR1* nucleosome association to PU.1 without (**b**) and with scFv (**c**). **d** to **f**, EMSA of *CX3CR1* nucleosome binding by ^DBD^C/EBPα (**d**), ^MBP-DBD^C/EBPα (**e**) and ^MBP-DBD^C/EBPα with scFv (**f**). For the experiments, the nucleosome concentration is 0.5 μM. The ratios of PU.1 proteins over the nucleosomes are 0, 0.25, 0.5, 1.0, 1.5, 2.0 and 2.5, respectively. The ratios of the C/EBPα proteins over the nucleosomes are 0, 1.0, 2.0, 3.0, 4.0, 6.0 and 8.0, respectively. **g**, Quantification of the *CX3CR1* nucleosome binding by PU.1 with and without scFv. **h**, Quantification of the *CX3CR1* nucleosome binding by ^MBP-DBD^C/EBPα with and without scFv. The K_d_^app^ values were obtained by fitting the data to Hill equation. At least 3 parallel experiments were performed for each assay. The error bars represent standard deviations.

**Extended Data Fig. 2.**
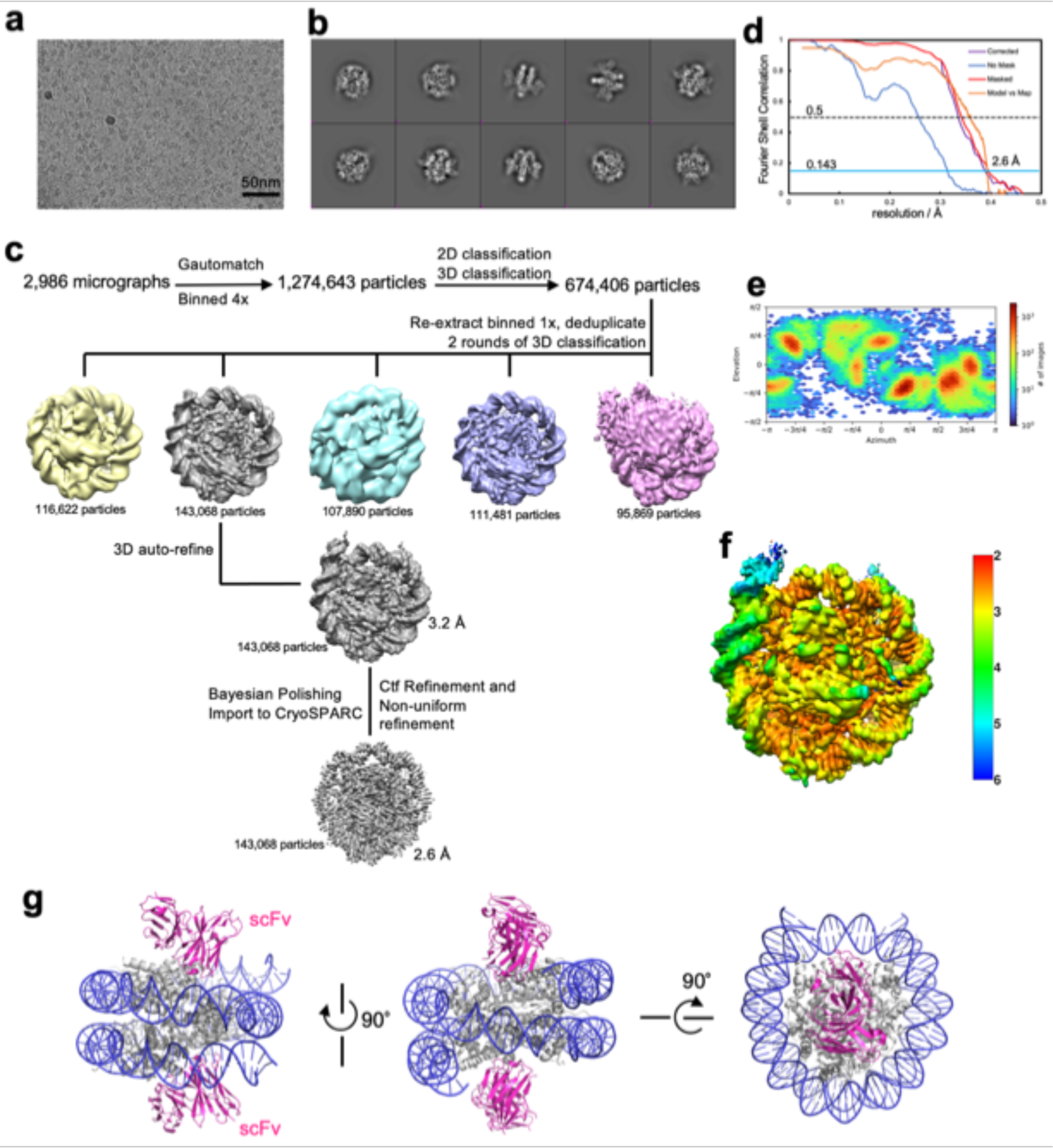
| Data collection and processing for the free 162 bp *CX3CR1* nucleosome. **a**, Raw image. The scale bar represents 50 nm. **b**, 2D classification results. **c**, Typical particle analysis. Note that resolution of the blue class can reach 2.9 Å and it has the same dyad location as the grey class. The dyad location of the other classes could not be determined due to lower resolution. **d**, FSC curves. **e**, Particle orientation distribution. **f**, Local resolution. **g**, Model of the nucleosome with scFv shows that scFv does not interact with DNA.

**Extended Data Fig. 3.**
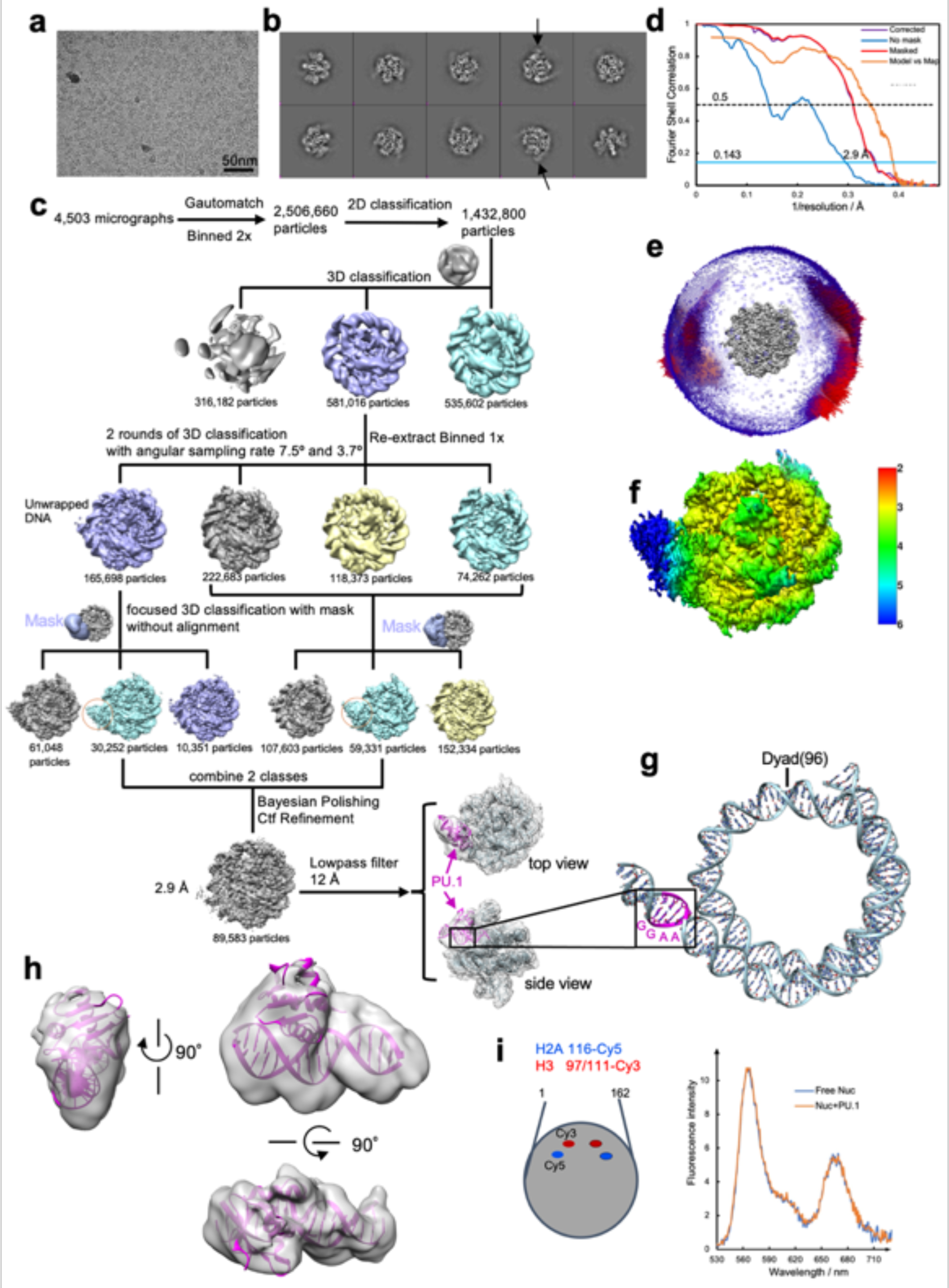
| Data collection and processing for the 162 bp *CX3CR1* nucleosome bound to PU.1. **a**, Raw image. The scale bar represents 50 nm. **b**, 2D classification results. The black arrow indicates the PU.1 features. **c**, Particle analysis. **d**, FSC curves. **e**, Particle orientation distribution. **f**, Local resolution. DNA was built from scratch based on the Cryo-EM map. At the nucleosome DNA exit site, there is a ‘GGAA’ located at the position with the extra density, indicating the PU.1 binding on the canonical site. **h,** Fitting of PU.1 crystal structure (PDB: 1PUE) into the extra density on the unwrapped DNA. **i,** FRET results show that H2A-H2B is not dissociated from the nucleosome core after PU.1 binding.

**Extended Data Fig. 4.**
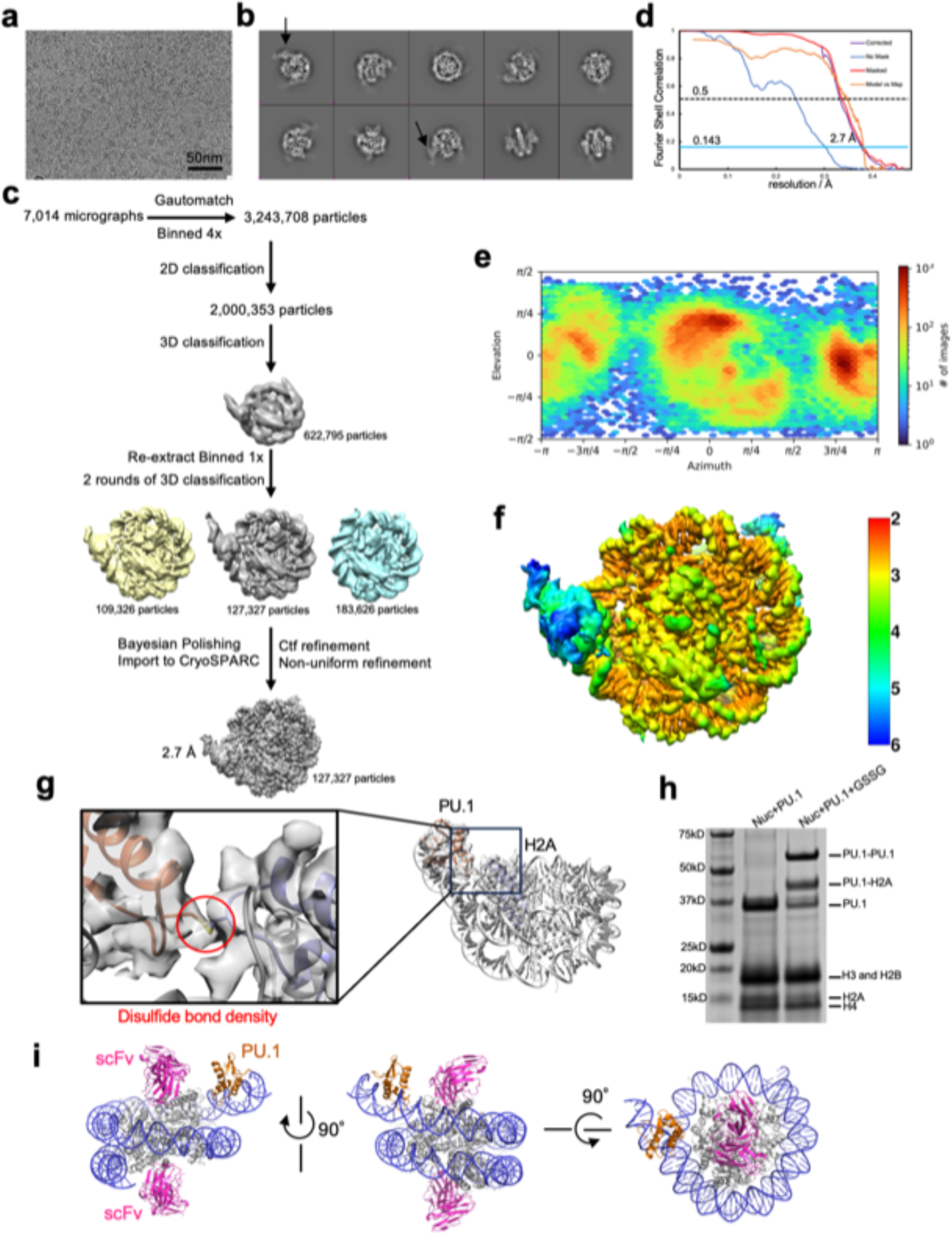
| Data collection and processing for the 167 bp *CX3CR1* nucleosome containing ^T77C^H2A bound to ^G220C^PU.1. **a**, Raw image. The scale bar represents 50 nm. **b**, 2D classification results. The black arrows indicate the densities of PU.1. **c**, Particle analysis. **d**, FSC curves. **e**, Particle orientation distribution. **f**, Local resolution. **g**, Cryo-EM map showing the disulfide bond between ^G220C^PU.1 and ^T77C^H2A. **h,** Non-reducing SDS-PAGE of nucleosome-PU.1 complex without and with the oxidant. **i,** Structure of the nucleosome-PU.1 complex with scFv shows that scFv does not interact with PU.1.

**Extended Data Fig. 5.**
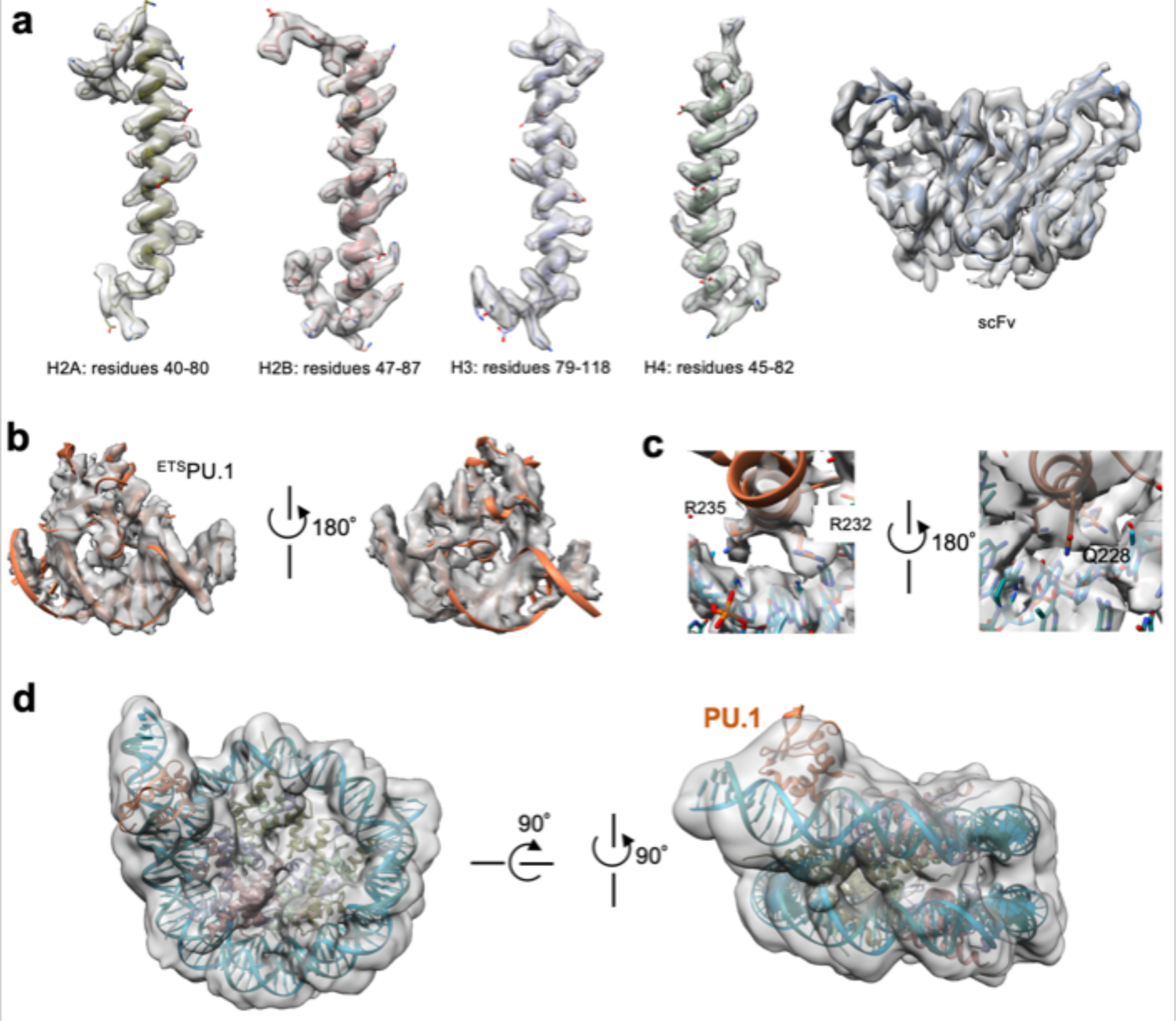
| Representative Cryo-EM densities and fitting of the structural models. **a,** Histones and scFv in the ^H2A-T77C^nucleosome-^G220C^PU.1-scFv complex. **b**, Fitting of the ^H2A-T77C^nucleosome-^G220C^PU.1 model to the wild-type nucleosome-PU.1 density map, showing that the disulfide bond does not change the complex structure. **c,** Illustration of the density map at the interface of ^ETS^PU.1 and DNA with highlights on the residues that interact with DNA. **d,** Fitting of ^H2A-T77C^nucleosome-^G220C^PU.1 structure into the density of the wild type nucleosome-PU.1-scFv complex.

**Extended Data Fig. 6.**
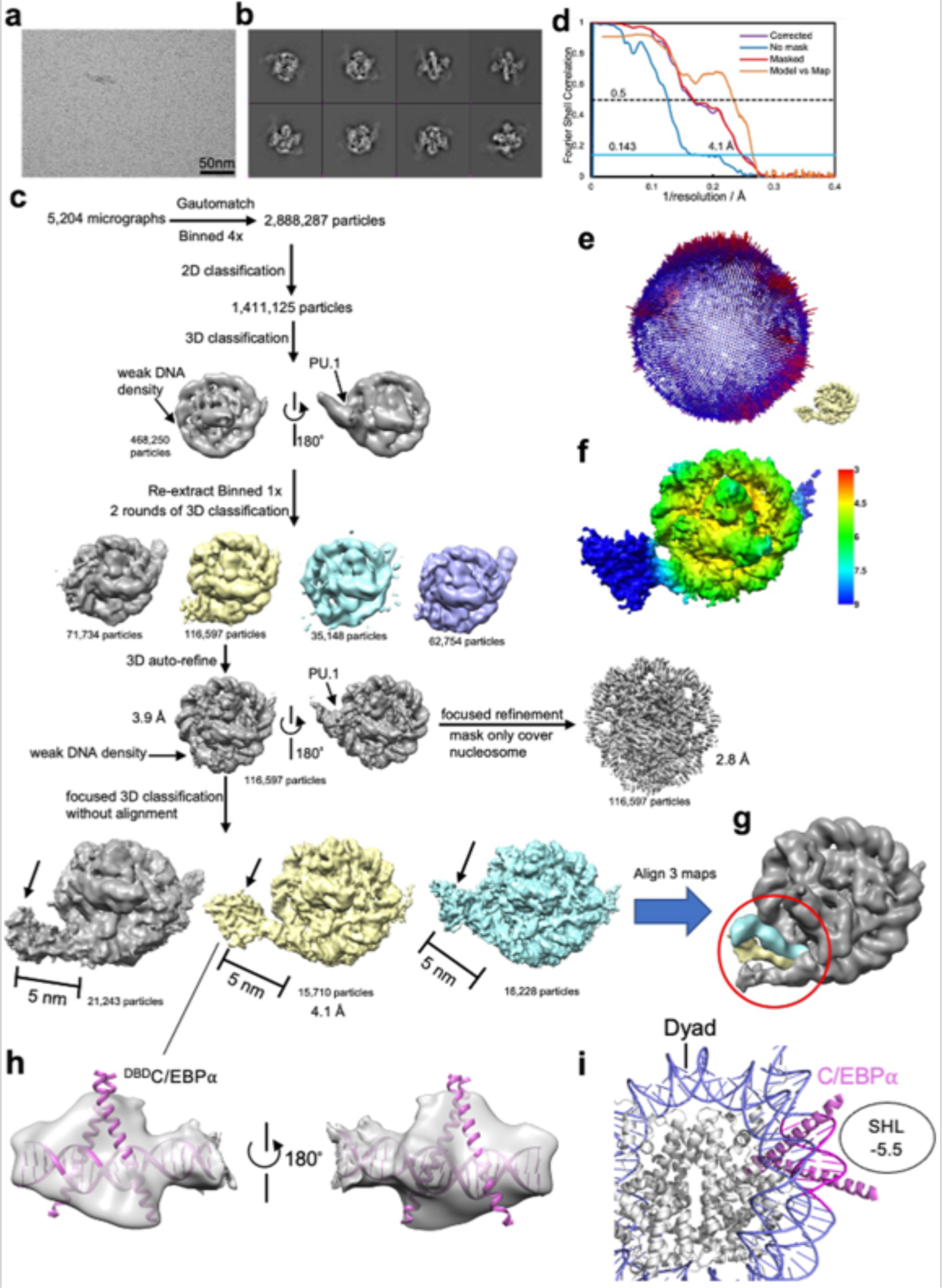
| Data collection and processing for the 167 bp *CX3CR1* nucleosome containing ^T77C^H2A bound to ^G220C^PU.1 and ^MBP-DBD^C/EBPα. **a**, Raw image. The scale bar represents for 50 nm. **b**, 2D classification results. **c**, Particle analysis. **d**, FSC curves. **e**, Particle orientation distribution. **f**, Local resolution. **g**, Alignment of three classes with different entry site DNA unwrapping degrees. **h**, Fitting of C/EBPα crystal structure (PDB: 1NWQ) into the extra density at unwrapped entry site DNA. **i,** Modeling of binding of C/EBPα binding to the fully wrapped nucleosome at the location corresponding to that in **g**.

**Extended Data Fig. 7.**
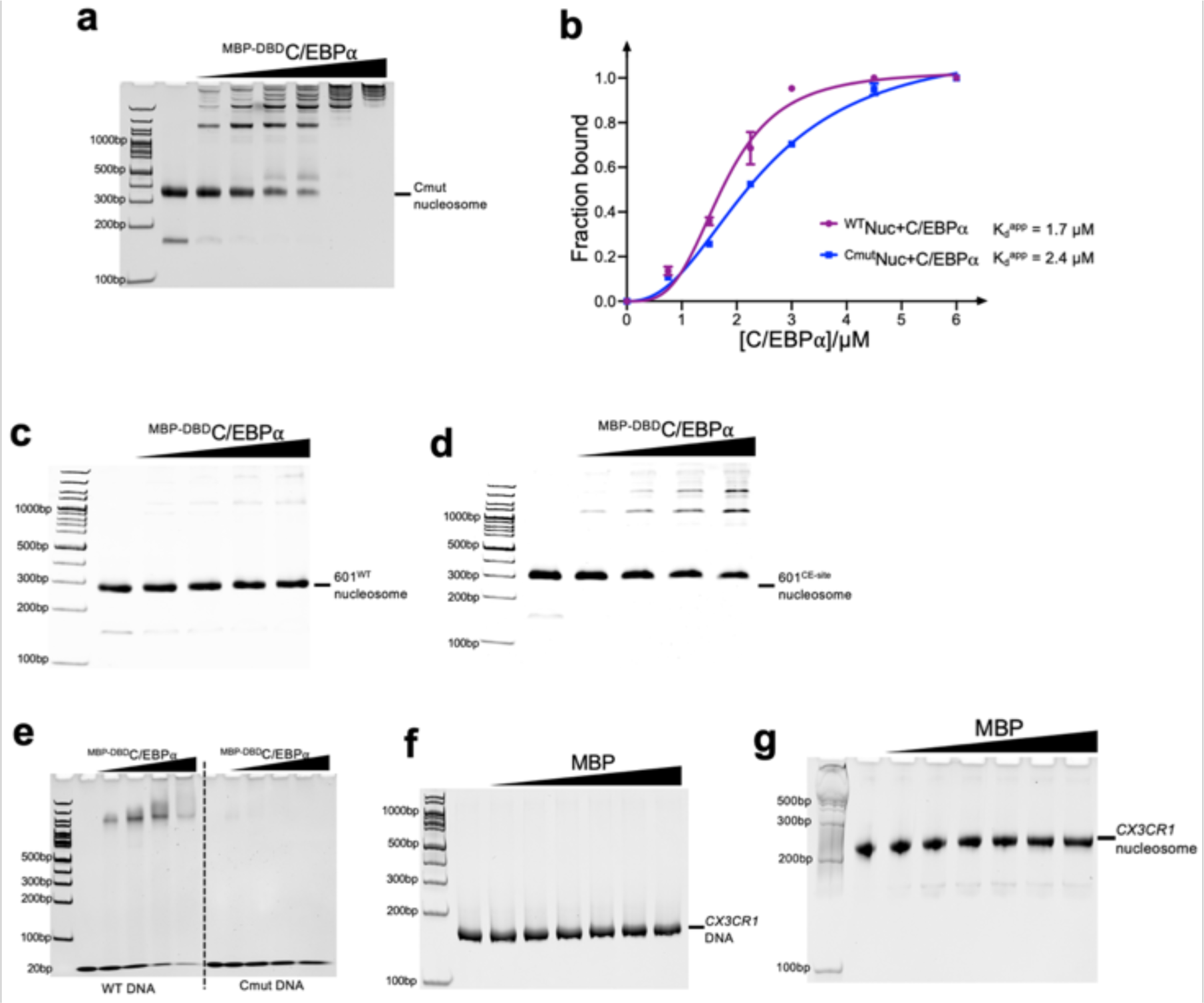
| Verification of C/EBPα binding location. **a,** EMSA of ^Cmut^nucleosome (the C/EBPα binding location ‘CAGC<u>TGG</u>TTG’ suggested by the cryo-EM result was mutated to ’CAGC<u>AAC</u>TTG’) binding by ^MBP-DBD^C/EBPα. For the experiment, the nucleosome concentration is 0.5 μM. The ratios of the C/EBPα proteins over the nucleosomes are 0, 1.0, 2.0, 3.0, 4.0, 6.0 and 8.0, respectively. **b,** Quantification of the wild type *CX3CR1* nucleosome (Extended Data Fig. 1**h**) and ^Cmut^nucleosome binding by ^MBP-DBD^C/EBPα. The apparent K_d_^app^ values were obtained by fitting the Data to Hill equation. At least 3 parallel experiments were performed for each assay. The error bars represent standard deviations. **c** and **d,** EMSA of the W601 nucleosome (**c**) and 601^CE-site^ nucleosome (‘CAGCTGGTTG’ was inserted into W601 at the corresponding location in *CX3CR1* nucleosome, illustrated by the diagram on the right) (**d**) binding by ^MBP-DBD^C/EBPα. For the experiment, the nucleosome concentration is 0.5 μM. The ratios of the C/EBPα proteins over the nucleosomes are 0, 1.0, 2.0, 3.0, 4.0, respectively. **e,** EMSA of 20 bp DNA containing wild type C/EBPα binding site (5’-ATCTT<u>CAGC**TGG**TTG</u>CTGAG-3’) and Cmut site (5’-ATCTT<u>CAG**CAA**CTTG</u>CTGAG-3’). For the experiment, the DNA concentration is 5 μM. The ratios of C/EBPα proteins over the DNA are 0, 0.5, 1.0, 1.5 and 2.0, respectively. **f** and **g,** EMSA of *CX3CR1* DNA (**f**) and nucleosome (**g**) binding by MBP. For the experiment, the DNA and nucleosome concentrations are all 0.5 μM. The ratios of the MBP protein over the DNA and nucleosome are 0, 1.0, 2.0, 3.0, 4.0, 6.0 and 8.0, respectively.

**Extended Data Fig. 8.**
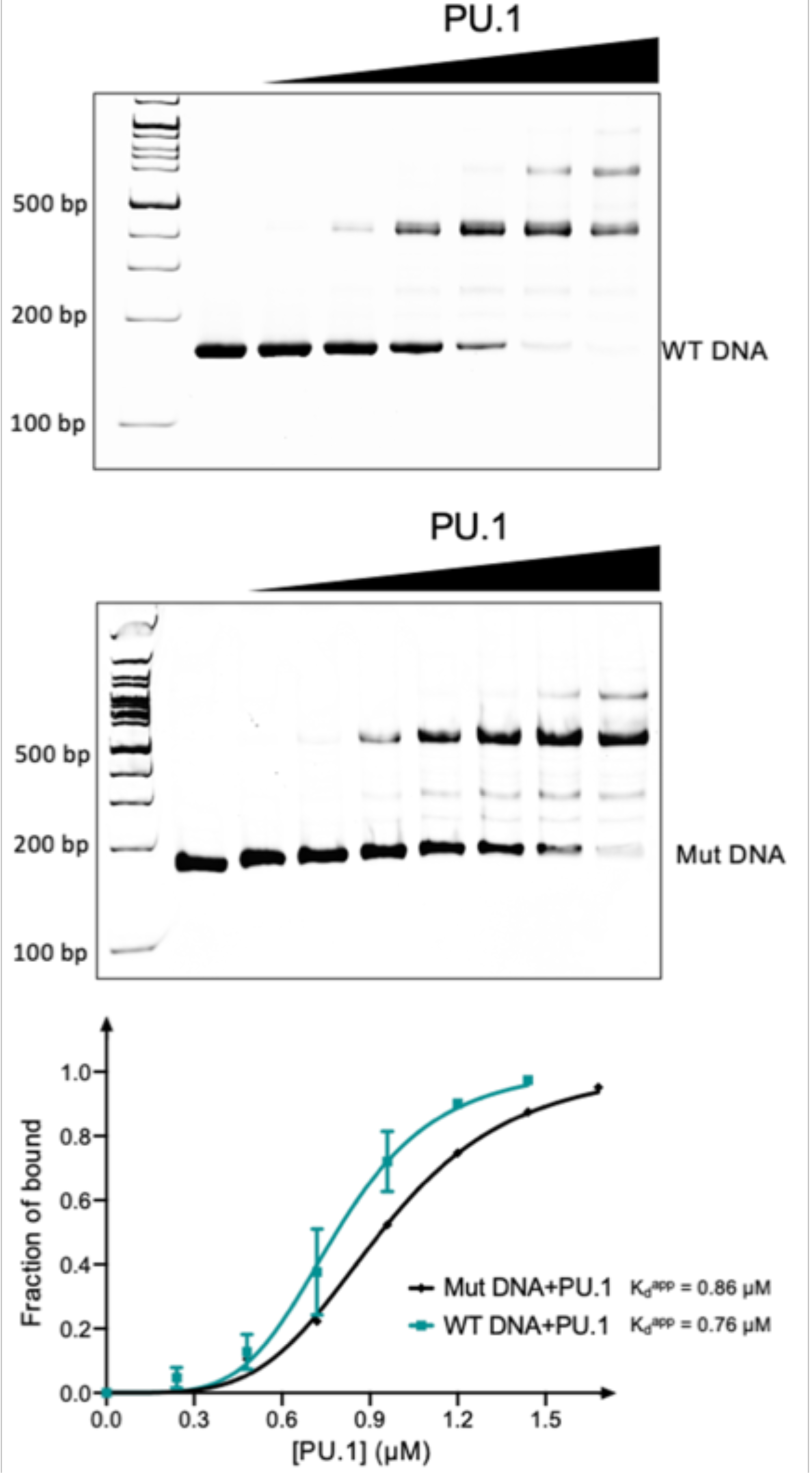
| Interactions between PU.1 and free *CX3CR1* DNA. EMSA of PU.1 and the 162 bp wild type *CX3CR1* and mutated (GG<u>AA</u> in the two sites near the dyad to GG<u>GG</u>) DNA (Figure 1. **a**). The DNA concentration is at 0.5 μM. The ratios of the PU.1 proteins over the WT DNA are 0, 0.25, 0.5, 1.0, 1.5, 2.0 and 2.5, respectively. The ratios of the PU.1 proteins over the mutated DNA are 0, 0.25, 0.5, 1.0, 1.5, 2.0, 2.5 and 3.0, respectively. The bottom panel shows the quantification of WT and mutated DNA binding by PU.1. The K_d_^app^ values were obtained by fitting the data (fraction of the nucleosome bound to PU.1 versus PU.1 concentration) to Hill equation. At least 3 parallel experiments were performed for each assay. The error bars represent standard deviations.

**Extended Data Fig. 9.**
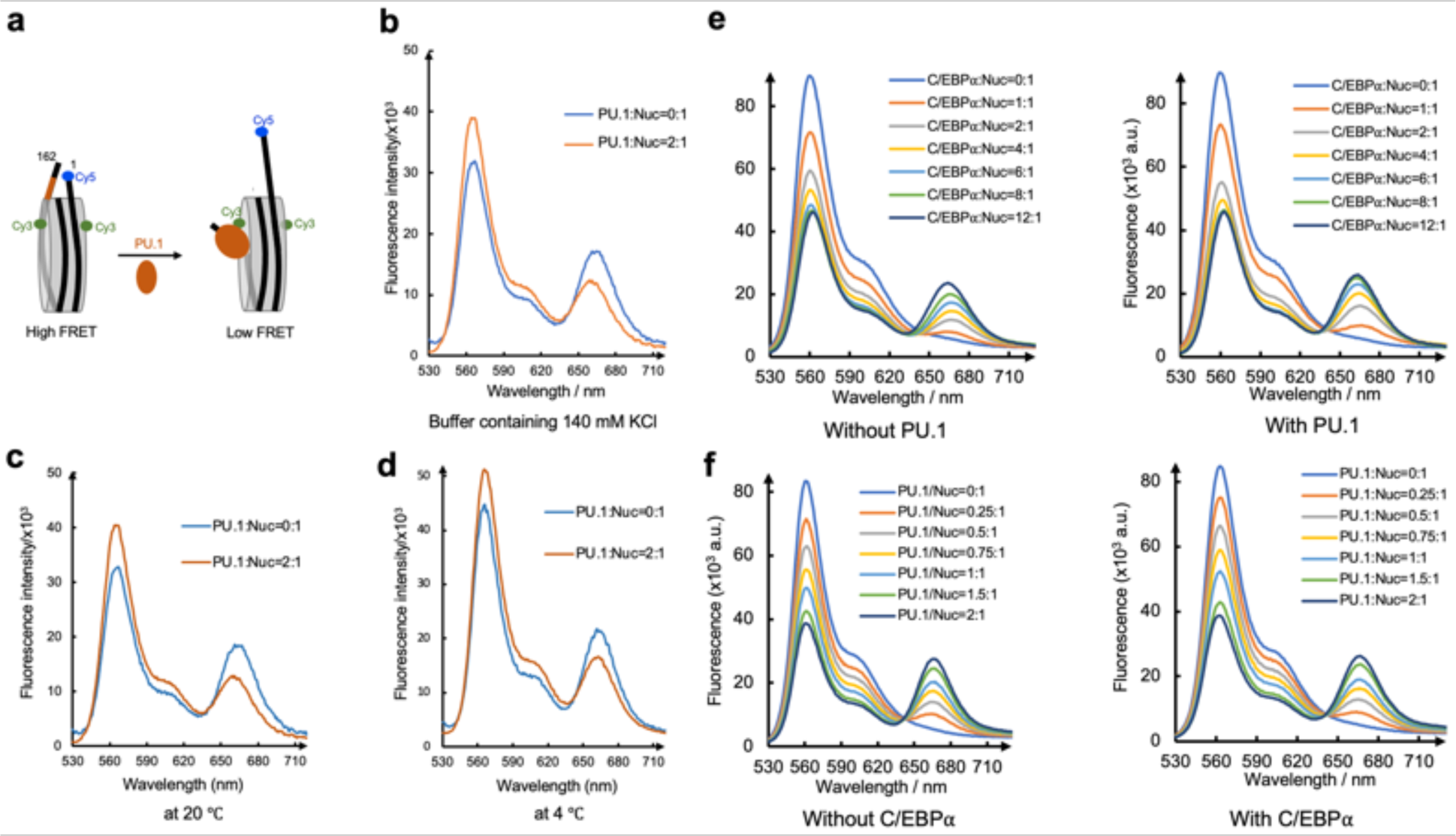
| The FRET assay confirms DNA repositioning and PU.1-facilitated nucleosome binding by C/EBPα.. **a,** Design of the FRET assay. Cy3 and Cy5 are labeled on the H2A residue 116 and the end of the DNA at the entry site. **b,** FRET in buffer containing 140 mM KCl. **c,** FRET at 20 ℃. **d,** FRET at 4 ℃. **e**, FRET assay with Cy3 labeled on DNA entry site (nucleotide 19) and Cy5 on C/EBPα in the absence (left panel) and presence (right panel) of PU.1. **f.** FRET assay with Cy3 labeled on DNA exit site and Cy5 on PU.1 in the absence (left panel) and presence (right panel) of C/EBPα.

**Extended Data Fig. 10.**
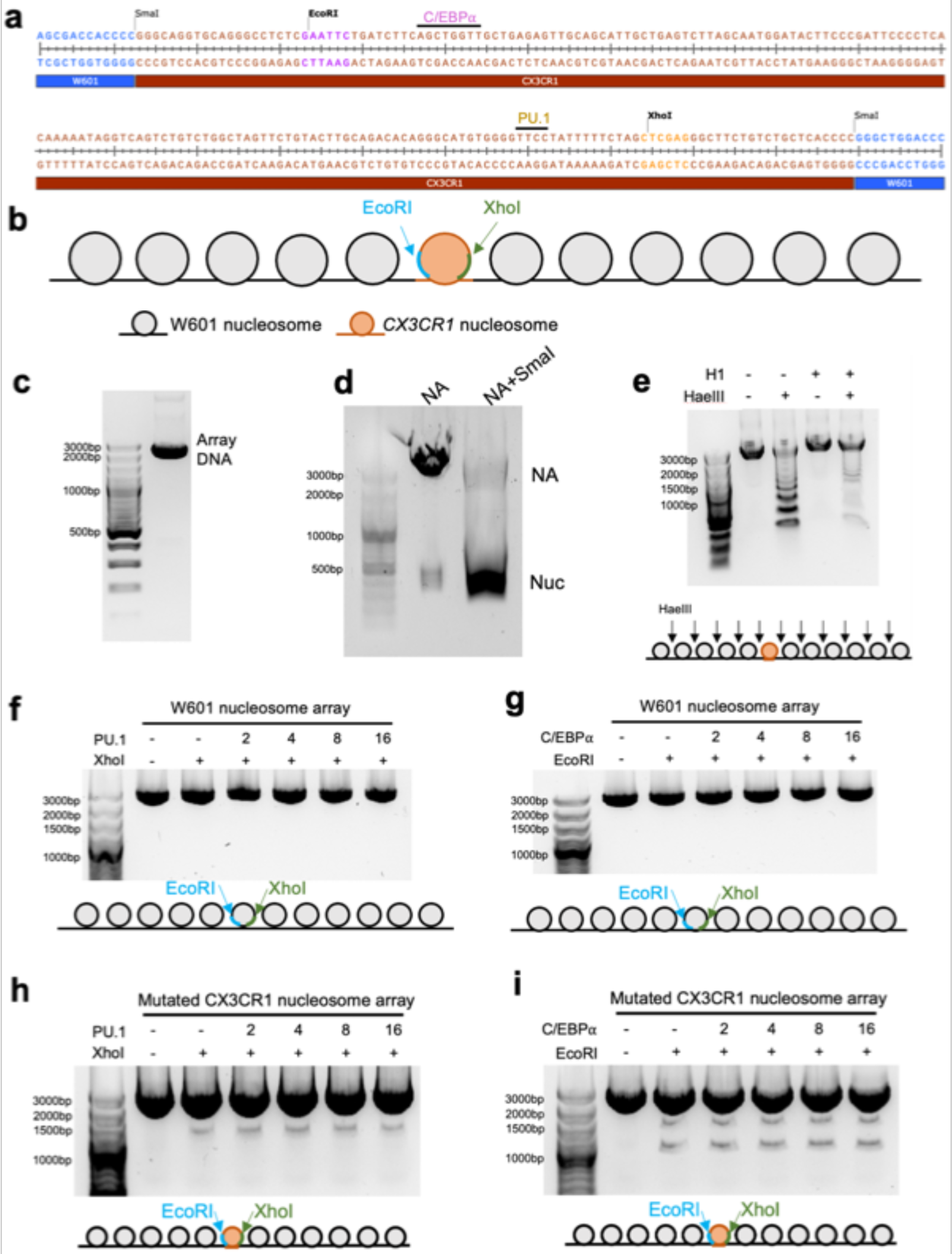
| PU.1 and C/EBPα facilitate endonuclease digestion of the *CX3CR1* chromatosomal DNA in the H1-condensed nucleosome array. **a**, Sequence and the endonuclease sites in the *CX3CR1* and neighboring Widom 601 (W601) DNA. **b.** The diagram shows the arrangement of the nucleosome array. **c**, Agarose gel showing purified 12 x 197 bp DNA. **d**, Agarose gel showing reconstituted nucleosome arrays digested by SmaI endonuclease. SmaI locates in all the linker DNA, so there are only mono-nucleosomes with 197bp DNA after cutting. **e,** Agarose gel showing reconstituted nucleosome arrays digested by HaeIII enzyme with and without H1.4. HaeIII exists in the linker DNA of both W601 and *CX3CR1* nucleosomes. **f** and **g**, Agarose gels showing the digestion of the H1.4-condensed nucleosome array by XhoI (**f**) and EcoRI (**g**) enzymes with different ratios of PU.1 and C/EBPα, respectively, over the W601 nucleosome array. **h** and **i**, Agarose gels showing the digestion of the H1.4-condensed PU.1 motif-mutated nucleosome array by XhoI (**h**) and the H1.4-condensed CEB/Pα motif-mutated nucleosome array by EcoRI (**i**) enzymes with different ratios of PU.1 and C/EBPα over the mutated nucleosome array, respectively.

